# Intrinsic excitability controls structural plasticity of cerebellar climbing fibers

**DOI:** 10.1101/2025.11.24.690115

**Authors:** Mattia Musto, Matilde Bergamini, Alessandra La Terra, Antonella Marte, Giacomo Marenco, Fabrizio Loiacono, Fabio Benfenati, Giorgio Grasselli

**Affiliations:** Center for Synaptic Neuroscience and Technology (NSYN), Istituto Italiano di Tecnologia (IIT), Genoa, Italy; IRCCS Ospedale Policlinico San Martino, Genoa, Italy; Department of Experimental Medicine (DIMES), University of Genoa, Genoa, Italy; Department of Neurosciences, Rehabilitation, Ophthalmology, Genetics, Maternal and Child Health (DINOGMI), University of Genoa, Genoa, Italy; Department of Pharmacy (DIFAR), University of Genoa, Genoa, Italy

## Abstract

In the cerebellum, climbing fibers (CFs) convey a teaching signal to Purkinje cells (PCs) that is crucial for cerebellar learning. They undergo activity-dependent synaptic plasticity during development and learning, and their transverse branches have an activity-dependent control of mobility. The maintenance of their structure was shown to rely on the growth-associated protein 43 (GAP-43), which is highly expressed in these fibers under basal conditions in adults and which is controlled by intracellular calcium levels and phosphorylation. Although activity-dependent structural plasticity occurs in several axonal fibers throughout the brain, it remains unclear whether this is also the case for the main branches of CFs. Here, we investigate this issue using *in vivo* knock-down of voltage-gated sodium channels (NaVs) in the CF-generating neurons of the inferior olive, and analyzing CF three-dimensional morphology, density and function of synaptic contacts with PCs. We show that the decrease in intrinsic excitability causes a reduction in the number and length of CF branches and that similar effects are caused by the knockdown of GAP-43, suggesting that GAP-43 may mediate this response. We also show that the retraction of CF fibers is associated with a seemingly compensatory increase in the density of CF synaptic terminals and PC dendritic spines, specifically in the area where PCs receive CF inputs. These plastic modifications are associated with a decrease in paired-pulse ratio and an increase in the kinetics and transferred charge of synaptic currents. Our data show that CFs can undergo activity-dependent structural plasticity changes affecting synaptic transmission and suggest the possibility that they contribute to encoding memories in the cerebellum and, if disrupted, to the pathophysiology of cerebellar diseases.

## INTRODUCTION

Increasing evidence has shown that memories are encoded in neuronal circuits by multiple and complementary forms of activity-dependent plasticity occurring at several sites, and not only by changes in synaptic strength (De Zeeuw et al., 2021; Gao et al., 2012; Holtmaat & Caroni, 2016). Several non-synaptic forms of plasticity have been found to contribute to encoding memories, making the picture more complex, and partially overcoming synaptic plasticity as the only paradigm of cellular mechanism underlying learning (Titley et al., 2017). These include forms of non-synaptic activity-dependent changes involving intrinsic excitability (i.e., intrinsic plasticity) (Daou & Margoliash, 2021; Debanne et al., 2019; Grasselli et al., 2020; Ohtsuki et al., 2020; Zhang & Linden, 2003), firing patterns (e.g., pause plasticity in the cerebellum; (De Zeeuw, 2021; De Zeeuw et al., 2021; Grasselli et al., 2016), and the morphology of cellular elements (i.e., structural plasticity; (Holtmaat & Caroni, 2016)). All these mechanisms are likely able to contribute to forming engrams (Ryan et al., 2021). However, the diversity of cellular players and events in the brain suggests that we may still be missing crucial pieces of such an intricate puzzle.

The cerebellum represents a well-established model to study cellular mechanisms underlying brain function and learning, due to the modular organization of the cerebellar cortex and to the relative simplicity of cerebellar motor learning paradigms (Ito, 2006). This has allowed us to identify cellular and molecular events mechanistically underlying the link between sensory inputs and behavioral (motor) output in the mammalian brain (De Zeeuw et al., 2021; Gao et al., 2012). This mechanistic role was shown for the long-term depression (LTD) of the excitatory synapses formed by parallel fibers (PFs) on Purkinje cells (PCs), which can be induced by protocols of paired stimulation of PFs together with the only other excitatory input of PCs, represented by climbing fibers (CFs) innervating PCs, usually in a one-to-one relation in mice (Busch & Hansel, 2023; Ito, 2001).

Within the olivo-cerebellar circuit, CFs play a crucial role in encoding information and driving synaptic plasticity in the cerebellar cortex: they are characterized by a periodic spontaneous tonic firing, as well as sensory-evoked firing of increased probability and frequency (De Zeeuw, 2021; De Zeeuw et al., 2021; Negrello et al., 2019). This CF-evoked firing is believed to represent a teaching signal for PF synapses on PCs, by dictating the induction of LTD at those PF synapses that fire in synchrony with the innervating CF (Gao et al., 2012; Otis et al., 2012). However, CFs can undergo LTD themselves, losing -in this way-the ability to induce LTD at PF-to-PC synapses (Coesmans et al., 2004). Modifications of CFs function can therefore cause changes in the rules of cerebellar plasticity (metaplasticity).

Climbing fibers also represent a timing signal for PCs, evoking a strong depolarization (the complex spike) followed by a delay (“CF-pause”) in the tonically generated action potentials (Jin et al., 2017), setting the timing of PC tonic firing patterns (Llinás, 2011; Person & Raman, 2011). The duration of this pause can undergo activity-dependent changes as well (Grasselli et al., 2016), which in turn modifies PC firing patterns.

The crucial role played by CFs in cerebellar physiology is further emphasized by the observation that alterations of the proper formation of CF synapses has been associated with several neuropathologies, such as spinocerebellar ataxias (Kuo et al., 2017; Louis et al., 2023), essential tremor (Kakei et al., 2021; Lang & Handforth, 2022; C. Y. Lin et al., 2014; Pan et al., 2020; Y. Wu & Louis, 2021), autism spectrum disorder (Piochon et al., 2014; Simmons et al., 2022) and schizophrenia (Veleanu et al., 2022).

Climbing fibers, initially considered to produce an invariable all-or-none synaptic response, evoke synaptic responses that can undergo activity-dependent synaptic plasticity (Hansel & Linden, 2000), and can be transiently modulated by sensory inputs (Gaffield et al., 2018, 2019; Najafi & Medina, 2013; Rowan et al., 2018; Yang & Lisberger, 2014; Zang & De Schutter, 2019). Some studies also showed that the morphology of CF branching (Bravin et al., 1999) or their a varicosities (Cesa et al., 2007) can change after a general blockade of neuronal activity or glutamatergic transmission in the cerebellar cortex (Bravin et al., 1999), and that the highly motile transverse branches of CFs (whose function is still largely debated) get stabilized if CF activity is pharmacologically increased (Nishiyama et al., 2007). However, these studies were based on systemic manipulations such as intraparenchymal or intraperitoneal infusion of drugs, leaving space to the possibility that complex circuit adaptations were underlying the observed CF effects. Activity-dependent changes in neuronal structure have been already observed in the cerebellum for synapses formed by mossy fibers filopodia and Golgi cells (Ruediger et al., 2011), PFs and PCs, mossy fiber collaterals and deep cerebellar nuclei (Boele et al., 2013; Kleim et al., 2002; Weeks et al., 2007), PC axon terminals and olivocerebellar axons (Foscarin et al., 2011), but not for those formed by CFs, at least during adulthood (Black et al., 1990; Connor et al., 2009; Kleim et al., 1998; Stevenson et al., 2021), despite activity-dependent plasticity of CFs has been shown to play a crucial role in the postnatal development of PC mono-innervation (Kano et al., 2018; Ohtsuki & Hirano, 2008)

Here, we investigate activity-dependence of CF morphology by chronically modulating their intrinsic excitability *in vivo* in juvenile mice, demonstrating that CFs undergo activity-dependent structural plasticity, involving the number and length of their branching, with opposite compensatory modifications of the density of synaptic terminals and post-synaptic dendritic spines. We also show that these modifications affect synaptic transmission, suggesting that they may contribute to encoding memories and that their impairments may participate in the pathophysiology of disorders involving the cerebellum.

## MATERIALS AND METHODS

### 1. VIRAL VECTORS

Third-generation lentiviral vectors modified to express shRNA sequences for the silencing of desired target genes were generated and produced as previously described (Grasselli et al., 2011). Briefly, five candidate 19 nt core sequences were designed according to previously established rationale (Reynolds et al., 2004) to target the identical regions of NaV1.1 (Scn1a, variants 1-2: NM_001313997.1, NM_018733.2) and NaV1.2 (Scn2a, variants 1-3: NM_001099298.3, NM_001346679.2, NM_001346680.1; **Fig. S1a**), which have a high degree of identity and are expressed in all neurons of the central nervous system, being therefore major responsible of the generation and propagation of action potentials (Duménieu et al., 2017). The selected target regions incidentally were identical in NaV1.3 (Scn3a, variants 1-7) as well. Each shRNA was formed by the sense core sequence and its antisense, separated by a loop sequence (TTCAAGAGA), preceded by a tetranucleotide (CCCC) and followed by a stop sequence (TTTTTGGAA) (Szulc et al., 2006; Tiscornia et al., 2003). The restriction sequences for Mlu1 and Cla1 were added for cloning. The candidate shRNAs were chemically synthetized, annealed and cloned into the transfer vector p207.pRRLsinPPTs.hCMV.GFP.WPRE (p207 for short, kindly provided by L. Naldini, Milan, Italy), previously modified with the insertion of a RNApolIII H1 promoter (upstream and divergent to the CMV promoter driving GFP expression) to make it suitable for shRNA expression (“p207-SH”). Low-titer preparations were tested on cerebellar primary cells prepared by real-time quantitative PCR for NaV1.2 expression as previously described (primers: CCTGGAAGGAAGTAGATTGACC, CCTCGGATGCTCAAGAGAGA; TaqMan Roche Universal ProbeLibrary #55; reference gene: ATP5b; (Grasselli et al., 2011; Mandolesi et al., 2009)) (**Fig. S1b**). One vector (shRNA2) was selected for medium-to-high silencing efficiency, for further testing at high-titer (core sequence: GAAGAAGTTTGGAGGTCAA targeting mouse Na 1.1, 1.2, and 1.3 homologous regions). The sequence of all vectors was confirmed by sequencing (IIT sequencing facility).

High-titer VSV-G-pseudotyped lentiviral particles were generated by calcium phosphate transfection in HEK293T cells with a mixture of 3 helper plasmids (encoding for the viral capside in pMDLg/pRRE #12251 and pRSV-Rev #12253 and the envelope in pMD2.VSVG #251049) and a transfer vector encoding GFP in combination with a shRNA sequence (specific for the target or a scramble control sequence). A similarly designed silencing vector targeting GAP-43 was previously published (shRNA core sequence: GAACATGCCTGAACTTTAA; (Grasselli et al., 2011)). The previously tested scramble sequence was used as a control (GACGAACGTATCCATATAT; (Grasselli et al., 2011)). Cells were cultured in Dulbecco’s modified Eagle’s medium (DMEM) supplemented with 10% fetal bovine serum (FBS), 1% GlutaMAX, 100 U/ml penicillin G, and 100 mg/ml streptomycin (all by ThermoFisher Scientific), and incubated at 37 °C and 5% CO_2_. At day zero, 2.7-3 × 10^6^ cells were plated in a 10 cm-dish. At day one, 24 h after plating, cells were transfected with 13 µg of pMLDg/pRRE, 3 µg of pRSV-Rev, 3.75 µg of pMD2.VsVG and 13 µg of transfer vector. The transfection mix containing DNA and calcium phosphate precipitates was washed out on day two, 14-16 h after transfection, with Dulbeccòs Phosphate-Buffered Saline (D-PBS, supplemented with 0.9 mM CaCl_2_ and 0.49 mM MgCl_2_ Mg^2+^), and re-incubated in 8-10 ml of fresh complete medium supplemented with Pyruvate. The resulting viral particles suspension was harvested 40-42 h after transfection, filtered through a 0.45-mm filter (Millipore Stericup), and concentrated by two ultracentrifugation steps of 1.5 h each at 4°C. The first centrifugation was performed at 26,000 rpm (∼115,000 rcf; Optima L-90K Ultracentrifuge; SW-32ti rotor, Beckman Coulter) with 4 mL of 20% sucrose solution at the bottom. The viral pellet obtained was then resuspended in DPBS (+Ca^2+^ and Mg^2+^) and ultraconcentrated with a second centrifugation at 23,000 rpm (∼56,700 rcf; MLS-50 rotor). The second viral pellet was eventually suspended in 20-40 µL DPBS (+ Ca^2+^ / Mg^2+^) and stored as 2 µL aliquots at -80° C until use. The functional titer of each preparation was estimated as follows: 100,000 HEK 293T cells/well were plated in a 6-well dish and were transduced 24 h later with serial dilutions of a frozen aliquot of the viral preparation (from 10^-3^ to 10^-7^), incubated for 72 h, rinsed in phosphate buffer solution (PBS), detached with 0.125% trypsin (1 min at RT), fixed with 4% PFA, resuspended in PBS and finally read with a flow-cytofluorimeter. The 2-3 dilutions obtaining ratios of GFP-positive cells in the range of 0.5-10% were used to estimate the viral titer based on the number of cells at the time of transduction (estimated 240,000/well) and the dilution factor, assuming that, at those low transduction frequencies, each GFP-positive cell was transduced by 1 viral particle. Only preparations with a viral titer of at least 1-3 × 10^9^ TU/ml (up to 5 × 10^10^) were used for *in vivo* injections.

### 2. ANIMALS

All animal experiments were designed in accordance with the national and European regulations for the care and use of experimental animals (Italian DL 26/2014; European Directive 2010/63/EU) and approved by the Animal Welfare Committee (*Organismo preposto al benessere degli animali*, OPBA) of San Martino Hospital and by the Italian Ministry of Health (Aut. 22418.120 145/2020-PR, 22418.N.LOU and 22418.N.CVI). Mice were housed in transparent cages with nesting material, food and water *ad libitum*, in 12:12 h light-dark cycles, together with littermates until surgery, and individually after surgery, with visual, auditory and olfactory access to littermates and other mice.

### 3. PRIMARY CELL CULTURES

High-density cortical neurons were prepared from C57BL/6J for protein extraction and electrophysiology experiments. Animals were sacrificed by CO_2_ inhalation, and embryos, at embryonic day 17-18, were immediately removed by cesarean section. Cerebral cortices were dissected by enzymatic digestion in 0.125% trypsin for 30 min at 37 °C and dissociated with a fire-polished Pasteur pipette. Primary cultures of dissociated cortical neurons were subsequently plated in poly-L-lysine-coated 35-mm multi-well plates (5 x 10^5^ cells/well) for protein extracts and on poly-L-lysine-coated 18-mm glass coverslips (4 x 10^4^ cells/well) for electrophysiology recordings. Neurons were maintained in a culture medium consisting of Neurobasal (Gibco) supplemented with B27 (2%, Gibco), Glutamax (1%, Gibco), penicillin-streptomycin (1%, Gibco) and kept at 37 °C in a 5% CO_2_ humidified atmosphere. After 7 days *in vitro* (DIV), neurons were transduced with lentiviral particles at 35 multiplicity of infection (MOI) by changing half of the medium.

### 4. WESTERN-BLOT

For Western blot analysis, total cell lysates were obtained from cultured primary cortical neurons, 7 days after viral transduction (14 DIV), suspended in lysis buffer (150 mM NaCl, 50 mM Tris-HCl pH 7.4, 1 mM EDTA pH 8, 1% TritonX-100; 100 ul/well) supplemented with a protease inhibitor cocktail (Cell Signaling). After 10 min of incubation on ice, cell lysates were collected and clarified by centrifugation (10 min at 10,000 x g at 4 °C). Protein concentration was determined using the BCA or Bradford assay (Pierce, Life Technologies). Equivalent amounts of protein were subjected to SDS-PAGE on Mini-PROTEAN® TGX™ Precast Protein Gels (Biorad) and blotted onto nitrocellulose membranes (Whatman). Blotted membranes were blocked for 1 h in 5% non-fat milk in Tris-buffered saline (10 mM Tris, 150 mM NaCl, pH 8.0) plus 0.1% Triton X-100 and incubated overnight at 4 °C with the following primary antibodies: anti pan-Nav (1:300; Sigma-Aldrich S8809, Merck), anti-GFP (1:2000; Invitrogen A-11122), anti-GAPDH (1:1000; Santa Cruz Biotechnology FL-335, sc-25778). Membranes were finally washed and incubated at room temperature (RT) for 1 h with horseradish peroxidase-conjugated secondary antibodies. Bands were revealed with the ECL chemiluminescence detection system (ThermoFisher Scientific).

### 5. STEREOTAXIC INJECTIONS

*In vivo* injections in the inferior olive (IO) were performed as previously described, with some adjustments (Grasselli et al., 2011). Juvenile (post-natal day 28-38) wild-type C57BL/6J mice (Charles River; mean age ± SD: P30 ± 2; mean weight ± SD: 15 ± 2g) were deeply anesthetized by isoflurane (1.9-2.5%) and placed in a stereotaxic apparatus on a heating pad with their nose-and ear-bars at the same height to minimize variables. Body temperature and respiratory rates were constantly monitored. Eye dehydration was prevented with artificial tears. The head and neck skin was shaved, disinfected, and incised. The dorsal neck muscles were retracted to expose the dura over the foramen magnum, and an opening was made to expose the brainstem. A borosilicate capillary (outer diameter: 1.0 mm; inner diameter: 0.58 mm) was pulled with a Sutter P-97 puller and manually broken to obtain micro-injection needles (outer diameter: 41-48 µm; inner diameter: 38-42 µm). These were mounted on a 10 µl glass syringe (Hamilton 701 RN), backfilled with mineral oil, then in an electronic injector (Quintessential Stereotaxic Injector, Stoelting). Micro-injection needles were front-filled with 1.5-2 µl viral suspension (thawed on ice and injected within 4-6 h). Injections were made unilaterally through the foramen magnum, just caudal to the bone, at 0.2 mm from the midline, with a 60-66° vertical angle and parallel to the sagittal plane, at a depth of 1.7-2.3 mm. A volume of viral suspension of ∼0.7 µl was delivered at a 50 nL/min rate, while the capillary was left in place for an additional 10 min before it was withdrawn. The incision was then sutured, and mice were treated with 0.25 mg of Diclofenac (16.7 mg / kg ca.) as anti-inflammatory and analgesic treatment, and 1 ml of 5 % glucose to rehydrate and facilitate recovery.

### 6. IMMUNOHISTOCHEMISTRY

#### Immunostaining for climbing fiber morphological analysis

Two weeks after viral injection, mice were deeply anesthetized with a ketamine-xylazine mix (5-10 mg/kg) and perfused through the heart left ventricle with ice-cold 4% paraformaldehyde (PFA) dissolved in 0.1 M PBS. The cerebellum and brainstem were isolated, post-fixed for 2 h at 4 °C in 4% PFA, and cryopreserved in a 30% sucrose solution at least overnight. Cerebellar sections were cut 30 µm thick with a freezing microtome, collected and stored in PBS for immediate use (within 5-7 days) or in anti-freezing solution (PBS 1X, 30% Ethylene glycol, 30% glycerol) for long-term storage at -20 °C. Slices were rinsed in PBS for at least 30 min from anti-freezing solution when necessary, permeabilized in 0.25% Triton-X-100 PBS (PBS-Tx) for 45 min, blocked for 1 h at RT with 10% normal goat serum (NGS) in PBS-Tx. They were then incubated overnight at 4°C with the following primary antibodies, diluted in blocking solution: mouse monoclonal anti-calbindin D-28k (1:1000; Swant 300) to stain PC cytosol; rabbit polyclonal anti-VGLUT2 (1:500; Synaptic System 135403) to stain CF synaptic terminals; rabbit polyclonal anti-VGAT (1:500; Synaptic System 131002) to stain inhibitory pre-synaptic terminals. Sections were rinsed with 0.25 % PBS-Tx for 3 times (for 30 min total), incubated for 2 h at RT with Alexa Fluor-conjugated secondary antibodies (anti-mouse Alexa Fluor 405 Invitrogen A31553; anti-rabbit Alexa Fluor 568 Invitrogen A11036), and then rinsed again with 0.25% PBS-Tx. Sections were mounted on microscope glass slides (Epredia Superfrost Plus Adhesion Microscope Slides, Fisher Scientific), air-dried, and cover-slipped using anti-fading mounting medium (Vectashield, Vector).

#### Immunostaining for dendritic spine density analysis

To obtain a higher tissue penetration of antibodies and efficiently stain dendritic spines, a 72 h permeabilization was performed with 0.25 % Triton X-100 solution at 4 °C. Sections were then incubated for 1 h with 0.5 % Tween in PBS at RT and then blocked for 1 h with a 10% NGS, 1% Bovine Serum Albumin (BSA) and 0.05 % Tween in PBS at RT. They were then incubated overnight at 4 °C with the following primary antibodies, diluted in blocking solution: mouse monoclonal anti-calbindin (1:200, Swant) to stain PC dendritic spines and rabbit polyclonal anti-VGLUT2 (1:500, Synaptic System) to stain CF pre-synaptic terminals. Sections were rinsed with 0.05 % Tween-PBS, then incubated for 2 h at RT with Alexa Fluor-conjugated secondary antibodies diluted in 0.05 % Tween-PBS (anti-mouse Alexa-fluor 568 and anti-rabbit Alexa-fluor 647, 1:100) and finally rinsed with 0.05 % Tween-PBS.

#### Immunostaining for the quantification of climbing fiber-specific protein expression levels

To avoid surface antigen masking of NaV channels caused by tissue fixative, the concentration of PFA was reduced in the perfusing solution to 0.5% and 0.5% of sucrose was added to equilibrate the osmolarity (C. Tian et al., 2014). Brainstems were included in OCT and rapidly frozen in dry ice before slicing (20 µm thick) with a cryostat. Sections were collected on glass slides. Before staining, they were rinsed in PBS for 30 min to remove OCT residues and permeabilized with a 0.3 % PBS-Tx at RT for 30 min, blocked for 1 h at RT with a 10% BSA in 0.3 % PBS-Tx solution. They were then incubated overnight at 4 °C with the following primary antibodies, diluted in a 10% BSA, 0.1% PBS-Tx solution: mouse monoclonal anti-panNaV (1:200; Sigma) to stain voltage-gated sodium channels, and rabbit polyclonal anti-calbindin (1:1000; Swant). Sections were then rinsed in 0.3 % PBS-Tx for five times (30 min total), incubated for 2 h at RT with Alexa Fluor-conjugated secondary antibodies (anti-mouse Alexa Fluor 568; anti-rabbit Alexa Fluor 647), and then rinsed again with 0.3% PBS-Tx. Sections were finally cover-slipped using anti-fading mounting medium (Vectashield, Vector).

### 7. MICROSCOPY AND IMAGE ANALYSIS

#### Confocal imaging

Images of immunolabelled samples were acquired with a Leica confocal imaging system (Leica TCR SP8) using a 63× oil immersion objective (1.4 N.A). For the analysis of CF morphology, z-stacks of images were acquired comprising GFP-expressing fibers in their integrity in PC layer and molecular layer (digital zoom: 1x; 2048×2048 pixel; 9.43 pixel/µm; z-step: 0.75 μm; 1 PAU). Only CFs with high level of GFP expression and not overlapping with other GFP-positive CFs were selected, to allow reliable 3D reconstruction.

For spine density analysis, to resolve dendritic spines and their synaptic inputs, images were acquired with higher resolution (digital zoom: 3×; 2048×1024 pixels; z-step: 0.2 μm; 33.28 pixel/µm; 0.7 PAU) and with improved signal-to-noise ratio (4 frame average). Only proximal dendrites (lateral diameter: 2.5-5.2 μm) were considered for the analysis.

For NaV expression analysis, z-stacks of images were acquired comprising GFP-expressing cell bodies in IO region (digital zoom: 1x; 2048×2048 pixel; 27.63 pixel/ µm; z-step: 0.25 μm; 1 PAU).

#### Quantitative confocal analysis

All images analyzed were processed with NIH Fiji software (Schindelin et al., 2012). The morphology of isolated GFP^+^ CFs was analyzed on images acquired from at least 3 animals per group, using always the same settings with only minor adjustments of laser power and photomultiplier gain, depending on fluorescence levels. Tracings from selected CFs were 3D reconstructed with the simple neurite tracer (SNT) plugin (Arshadi et al., 2021), to identify and measure transversal ramifications, total length of the fiber, and length of the PC dendrite innervated by the CF. Tracings were then analyzed by Sholl’s method (Sholl, 1953) by placing the center of analysis on CF segment located in the PC layer (radius step: 2 μm; radius span: 1 μm). The median number of intersections was calculated for each CF in the interval of interest, corresponding to the distance at which the maximum number of branches was present. The number of axonal varicosities, identified either morphologically or by VGlUT2 staining, was manually counted using Fiji.

The PC dendritic spine density was calculated by counting only the spines emerging from the lateral side of proximal or distal dendrites. Each identified spine was followed until it disappeared downstream and upstream in the image series, to exclude spines emerging from other dendritic segments (Mandolesi et al., 2009).

To quantify NaV expression, the mean intensity of the specific fluorescence signal associated with the anti-PanNa_V_ antibody was measured with Fiji on the maximum intensity projection of GFP^+^ cells localized in the IO region. The mean fluorescence intensity value was expressed as a percentage of the control.

### 8. ELECTROPHYSIOLOGY

#### Primary cell electrophysiology

Whole-cell patch clamp recordings were performed on primary cortical neurons 7 days after viral transduction (14 DIV). Patch pipettes, prepared from borosilicate glass (World precision instrument, Sarasota, Florida), pulled and fire polished to a final resistance of 2.5-4 MΩ, were filled with the internal solution containing: 126 mM K-Gluconate, 4 mM NaCl, 1 mM MgSO_4_, 0.02 mM CaCl_2_, 0.1 mM BAPTA, 15 mM Glucose, 5 mM HEPES, 3 mM ATP, 0.1 GFP, with pH adjusted to 7.3 with 1 M KOH. For all the experiments, cells were maintained in standard extracellular Tyrode solution containing: 140 mM NaCl, 2 mM CaCl_2_, 1 mM MgCl_2_, 4 mM KCl, 10 mM glucose, and 10 mM HEPES (pH adjusted to 7.25-7.35 with NaOH), supplemented with 50 µM Picrotoxin and 10 µM 6-cyano-7-nitroquinoxaline-2,3-dione (CNQX) to block GABA_A_ receptors and AMPA receptors, respectively to block most of glutamatergic excitatory and GABAergic inhibitory transmission.

Cells were visualized with a BX51W1 microscope (Olympus) equipped with a 40X objective, with a CCD camera (ORCA-R2 C10600, Hamamatsu). Patch-clamp recordings were performed using the MultiClamp 700B Microelectrode Amplifier (Axon CNS). Current-clamp recordings of action potential (AP) firing activity were performed at a holding potential of −70 mV, and action potentials (APs) were induced by injection of 50 pA current steps lasting 500 ms.

To study the biophysical properties of the single AP more in depth, a plot of the time derivative of voltage (dV/dt) versus voltage was performed (phase-plane plot). The first AP elicited by the minimal current injection was used to obtain the plot, from which the maximum rising slope, the AP peak, the maximum repolarizing slope, and the V_threshold_ were extracted. The voltage threshold was defined as the first voltage value at which dV/dt exceeded 4mV/ms without decreasing. The slope of phase-plots at the AP threshold (“kink”) was calculated using linear regression of the first 10 data points of the rising phase starting from the V threshold point. The electrophysiological recordings were analyzed using OriginPro 8 (OriginLab Corp., Northampton, Ma,USA) software.

#### Slice electrophysiology

Two weeks after viral injection, C57BL/6J mice (age: P42-49) were decapitated under isofluorane anesthesia. Sagittal slices of the cerebellar vermis, used for the recordings of evoked excitatory post-synaptic currents (eEPSC), were cut 250 μm thick with a Leica vibratome (VT1200S) and ceramic blades in ice-cold artificial cerebrospinal fluid (ACSF) containing the following: 124 mM NaCl, 5mM KCl, 1.25 mM Na_2_HPO_4_, 2 mM MgSO_4_, 2 mM CaCl_2_, 26 mM NaHCO_3_, and 10 mM D-glucose, bubbled with 95% O_2_/5% CO_2_. The slices were kept at RT in ACSF for at least 1 h, then transferred to a submerged recording chamber superfused with ACSF supplemented with picrotoxin (100 μM) to block GABA_A_ receptors. Whole-cell patch-clamp recordings were performed at RT under visual control with differential interference contrast optics combined using an Olympus F-view II CCD camera and an Olympus LUMPLFLN 40× objective, mounted on an Olympus BX61WI microscope. Patch pipettes were prepared from borosilicate glass (World precision instrument, Sarasota, Florida), pulled and fire polished to a final resistance of 2.5–4 MΩ, and filled with internal saline containing: 100 mM CsMeSO_4_, 50 mM CsCl, 2 mM Na_2_ATP, 0.3 mM Na_3_GTP, 1 mM MgCl_2_, 0.2 mM EGTA, 4 mM QX-314 (Lidocaine N-ethyl bromide) and 10 mM HEPES, with pH adjusted to 7.25–7.35 with 1 M CsOH and osmolarity adjusted to 295–305 mmol/kg with sucrose. To visualize the patched cell, a red fluorescent dye (10 mM) was added to the internal. The CF-stimulation pipettes were filled with ACSF. Recordings were performed at a holding potential of 0 mV. CFs were stimulated with the minimum intensity to evoke an EPSC by placing the stimulating electrode in the granular layer close to the Purkinje cell body. The pairing of the recorded dye-filled Purkinje cell with a GFP^+^ CF was then verified morphologically at the fluorescence microscope at the end of the recording to include only recordings from PC clearly innervated by transduced CFs.

### 9. STATISTICAL ANALYSIS

Pair-wise comparisons were made by Student’s *t*-test, except for data obtained by electrophysiology, which were made by Mann-Whitney’s *U*-test. Comparisons between cells which were repeatedly tested (Sholl analysis of CFs, intrinsic excitability test of primary neurons, pair-pulse ratio tested at several inter-stimulus intervals) were made by ANOVA for repeated measures (RM-ANOVA). The number of CFs has been considered as the statistical unit, represented on figures by the numbers on histograms, and “n” represents the number of CFs. Means ± SEM and individual values were represented in graphs. Statistical significance was diaplayed as follows: *p<0.05, **p<0.01, ***p<0.001.

## RESULTS

### 1. Effects of GAP-43 on CF morphology

We have previously shown that the physiological expression of the growth-associated protein 43 (GAP-43) in cerebellar CF not only is necessary for axonal sprouting in reaction to traumatic conditions (such as the partial lesion of the IO or laser microdissection of single CF branches), but also to sustain CF structure and stability (Allegra Mascaro et al., 2013; Grasselli et al., 2011), suggesting its possible role in mediating forms of activity-dependent structural plasticity. First, we assessed whether the effects of GAP-43 knockdown (GAP-43-KD), previously observed in Wistar rats and FVB mice (Grasselli et al., 2011), are also observed in mice of the C57BL/6J strain to increase the future transferability of the results and method to genetically modified mouse models (**Fig. 1a**). Moreover, we set up an advanced morphometric analysis in which CF reconstructions (including not only CF stalks but also CF tendrils innervating the same dendritic branches) were made 3D to increase the sensitivity of the measures (**Fig. 1b**). Under these conditions, we confirmed that GAP-43-KD causes a substantial reduction of the total length of the fibers (GAP-43-KD: 857.8±54.8 μm; control: 1506±90 μm, p<0.0001; **Fig. 1c,d**), of the length of the corresponding innervated dendritic territory (GAP-43-KD: 552.5±26.1 μm; control: 787.1±55.2 μm, p=0.0044; **Fig. 1d**), and of the number of CF branches (average at 40-120 μm from the PC: 4.27±0.09 in GAP-43-KD fibers, 6.78±0.23 in control fibers, p=3.7*10^-7^; vector x radius interaction: p=0.0010; F (89, 1335) = 1.554; **Fig. 1e,f**). We also confirmed that this fiber retraction resulted in a decrease of the total number of axonal varicosities (GAP-43-KD: 124±6.6; control: 208.5±13.0, p=0.0001; **Fig. 1g,h**). Differently from what previously reported, however, silencing GAP-43 did not affect the density of varicosities, either measured on the total length of CF branches (GAP-43-KD: 1.44±0.06 varicosities/10 μm; control: 1.36±0.07 varicosities/10 μm, p=0.44; **Fig. 1i**) or on the length of the innervated dendritic territory (GAP-43-KD: 2.25±0.08 varicosities/10 μm; control: 2.72±0.17, p=0,060; **Fig. 1l**). We additionally quantified the fraction of varicosities expressing the vesicular glutamate transporter VGLUT2, observing that it was higher after silencing GAP-43 (GAP-43-KD: 94.71±0.91 %; control: 80.8±3.73 %, p=0.0083; **Fig. 1m**), suggesting that the treatment may also affect CF synaptic transmission. Overall, these data confirm that the expression of GAP-43 (a calcium and PKC-regulated protein) (Denny, 2006) is necessary to maintain the structure of CF, suggesting an activity-dependent mechanism of control of CF structure.

**Figure 1.**
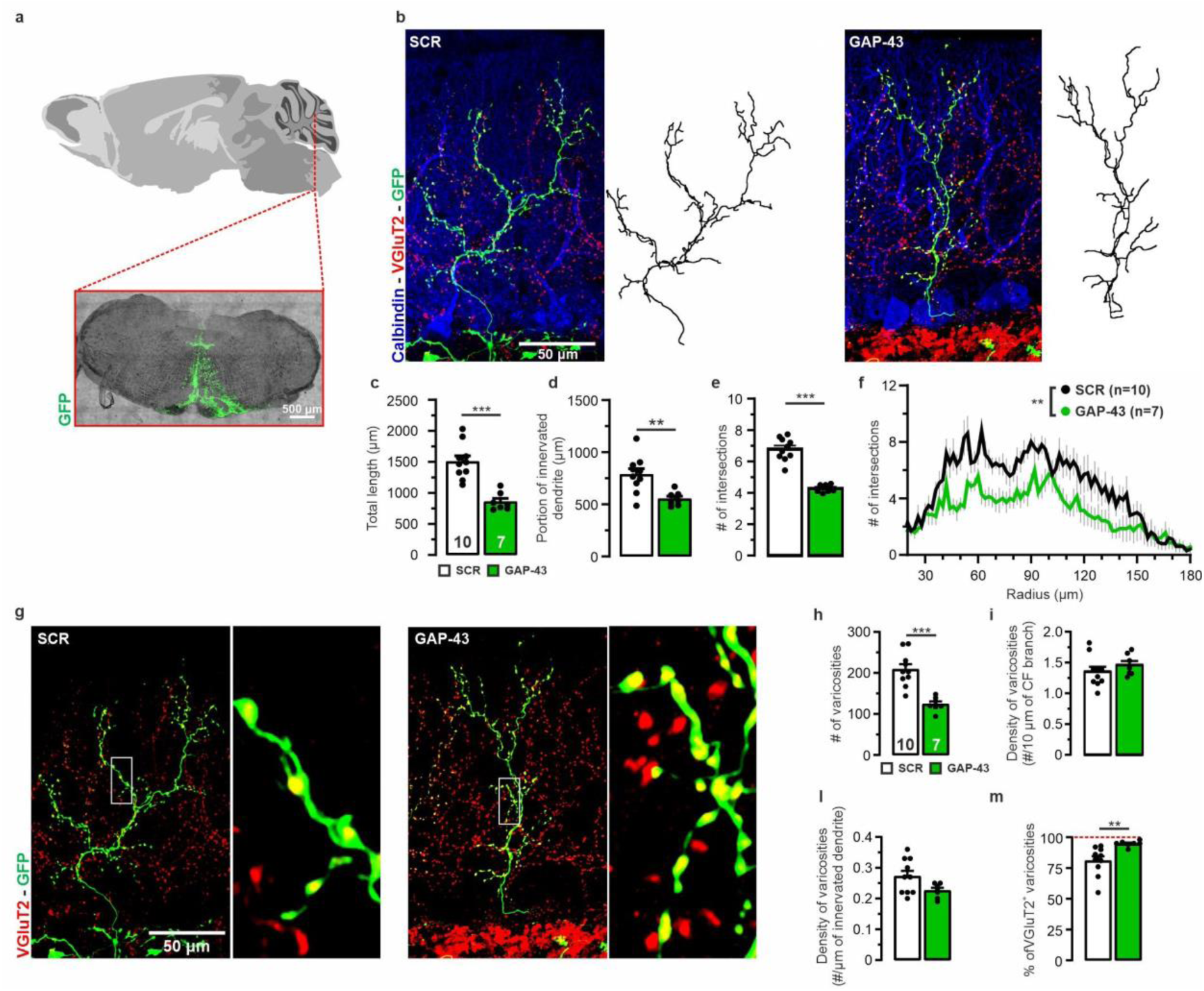
Knocking-down GAP-43 induces CF atrophy. **(a)** Injection site in the IO of GFP-expressing lentiviral vectors (brainstem coronal section): GFP (green) merged with brightfield. **(b)** Representative GFP-expressing CFs (green) in sagittal cerebellar sections immunostained for VGLUT2 (varicosities; red) and calbindin (PCs; blue), and corresponding 3D reconstructed traces (black) for control vectors (SCR*, left*) and after GAP-43-KD (*right*). **(c-f)** GAP-43-KD causes a reduction of the total length of CFs (**c**), of the portion of dendrite innervated by the CF (**d**), and of the number of CF branches (*e*), as assessed by Sholl analysis at 40-120 μm from the PC soma (**f**). **(g)** Representative GFP-expressing CFs immunostained for VGLUT2 (varicosities, in red; higher magnification of the images shown in b). **(h-l)** GAP-43-KD causes a decrease in the total number of varicosities (**h**) due to CF atrophy, accompanied by an unaffected density of varicosities over the length of the CF (**i**) and on the length of PC innervated dendrite (**l**). **(m)** The relative number of VGLUT2-expressing varicosities is higher after GAP-43-KD. Scale bars: 500 µm (**a**) and 50 µm (**b, g**). *p<0.05; **p<0.01; ***p<0.001.

### 2. Knock down of NaV channels in IO neurons

To assess the possibility of an activity-dependent control of CF morphology, we aimed at inducing a chronic reduction of CF activity. To do so, we used an shRNA sequence specific for the pore-forming subunits NaV1.1 and NaV1.2 of voltage-gated sodium channels (**Fig. S1**). These have a high degree of identity; they are widely expressed in the brain, including the cerebellum (Schaller & Caldwell, 2003), and play a major role in the generation and propagation of APs (Duménieu et al., 2017). Expression of the shRNA reduced the overall expression level of all NaV isoforms in primary cortical neurons to 29 ± 6% of the scrambled shRNA control (SCR) after 7 days of viral transduction, as assessed by Western blot with a pan-NaV antibody (p=0.0004; **Fig. 2a,b**). In parallel, we observed that the cortical neurons that were knocked down of NaV channels (NaV-KD) had a significant decrease in intrinsic excitability (measured as the number of APs evoked by the injection of a depolarizing current of increasing amplitude; KD x injected current interaction: F (19,266) = 2.394, p=0.0012; **Fig. 2c,d**), as well as a decrease in the peak amplitude (NaV-KD: 24.4±3.2 mV; control: 41.6±3.7, p=0.0079; **Fig. 2e**) and rising slope (NaV-KD: 75.2±13.1 mV/ms; control: 167.4±27.9 mV/ms, p=0.023; **Fig. 2f**) of their APs, under comparable recording conditions (input resistance and holding membrane potential; **Fig. 2g,h**).

**Figure 2.**
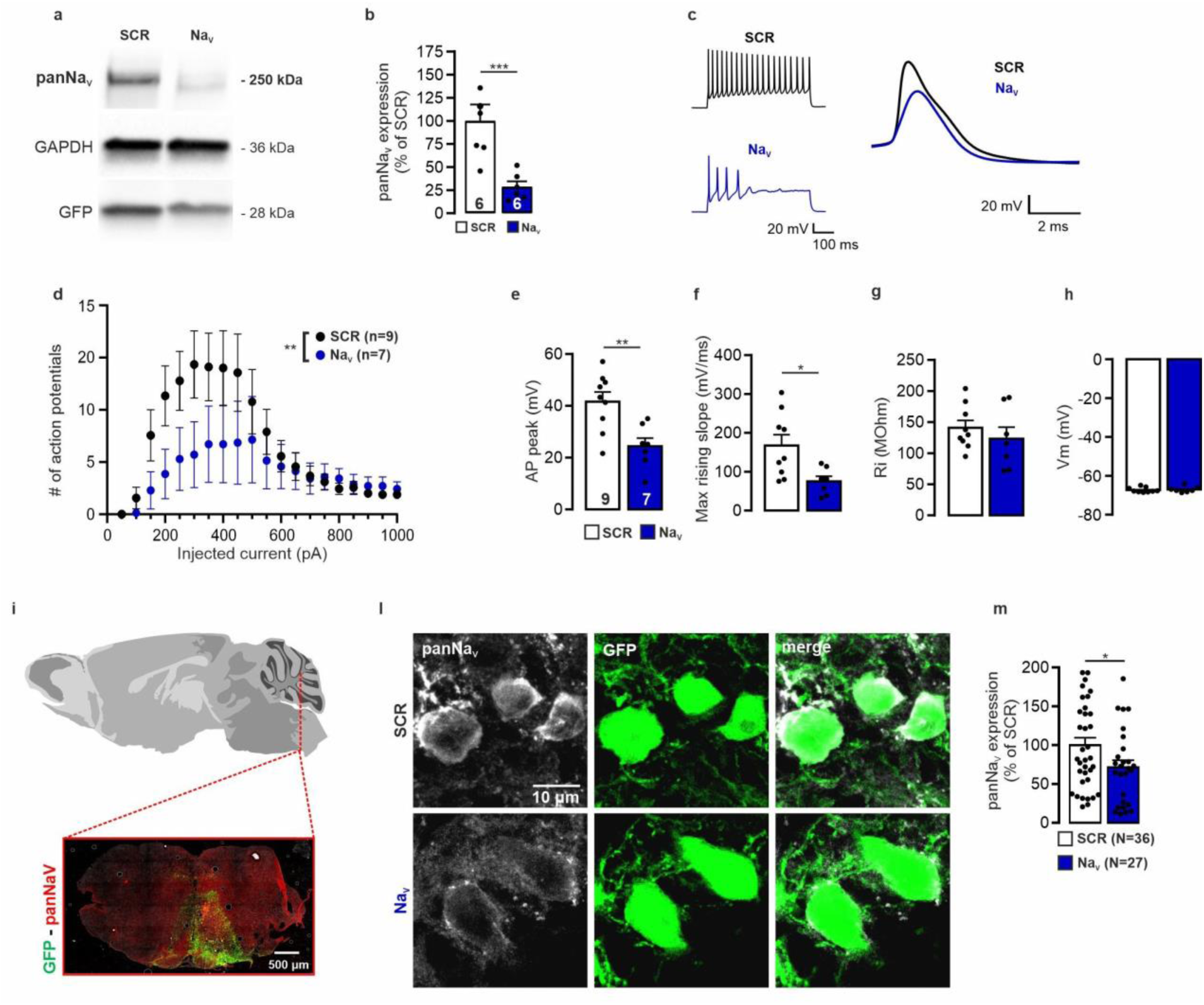
*In vivo* knock-down of NaV1.1/1.2 by sh-RNA-expressing lentiviral vectors. **(a,b)** *In vitro* efficacy of NaV1.1/1.2 knock-down (NaV-KD). Representative western blotting with panNaV antibody (a) and the respective quantification **(b)** 7 days after transduction of primary cortical neurons (14 DIV) with lentiviral particles expressing either NaV1.1/1.2 shRNAs or a scramble shRNA sequence (SCR). **(c)** Representative AP traces evoked by a square injection of depolarizing current (*left*) and corresponding detail of single APs (*right*), recorded in cortical neurons 7 days after transduction (14 DIV). **(d)** The number of evoked APs as a function of the injected current shows a decreased excitability after NaV-KD. **(e,f)** Analysis of AP kinetics: the peak (**e**) and the maximum rising slope (**f**) are slowed down by NaV-KD. (**g,h**) The intrinsic properties input resistance (**g**) and resting membrane potential (**h**) are unaffected by the treatment. **(i)** Injection site of GFP-expressing silencing vectors (green) shown on a coronal section of brainstem immunostained with panNaV antibodies (red). **(l)** GFP-positive cells (green) in the IO immunostained with panNaV antibodies (white; maximum projection of z-stack). **(m)** Quantification of panNaV immunoreactivity in GFP-positive areas of confocal z-stacks from samples treated with either NaV-KD or SCR vectors. Scale bars: 500 µm (**i**) and 10 µm (**l**). *p<0.05; **p<0.01; ***p<0.001.

When juvenile mice (P28-38) were transduced *in vivo* to target the IO, we observed a decrease in the overall expression of all NaV channel isoforms in GFP^+^ neurons already 2 weeks after the viral injection, indicated by a significantly lower immunoreactivity to the panNaV antibody (down to 71 ± 9% of the control; **Fig. 2i-m**; p=0.038), confirming the *in vivo* efficacy of the lentiviral NaV-KD vector.

### 3. Effect of the of NaV-KD on CF morphology

We then investigated, in the mice injected with the NaV-KD vectors in the IO, the effects of the downregulation of NaVs on CF morphology. We observed that, similarly to what observed for GAP-43-KD (see Fig. 1a-f), also NaV-KD causes a retraction of CF branches already two weeks after transduction (**Fig. 3a,b**), consisting in a reduction of the total length (NaV-KD: 1017±71 μm; control: 1506±90 μm; p=0.0003; **Fig. 3c**), of the length of innervated dendrite (NaV-KD: 583.5±36.4 μm; control: 787.1±55.2 μm; p=0.0043; **Fig. 3d**) and of the number of branches, as assessed by Sholl analysis (average within 40 and 120 μm from the PC soma: NaV-KD 4.69±0.4 intersections, control 6.78 ± 0.23 intersections, p=1.2*10^-4^; vector x radius interaction F (80, 1680) = 1.320, p=0.033; **Fig. 3e,f**).

**Figure 3.**
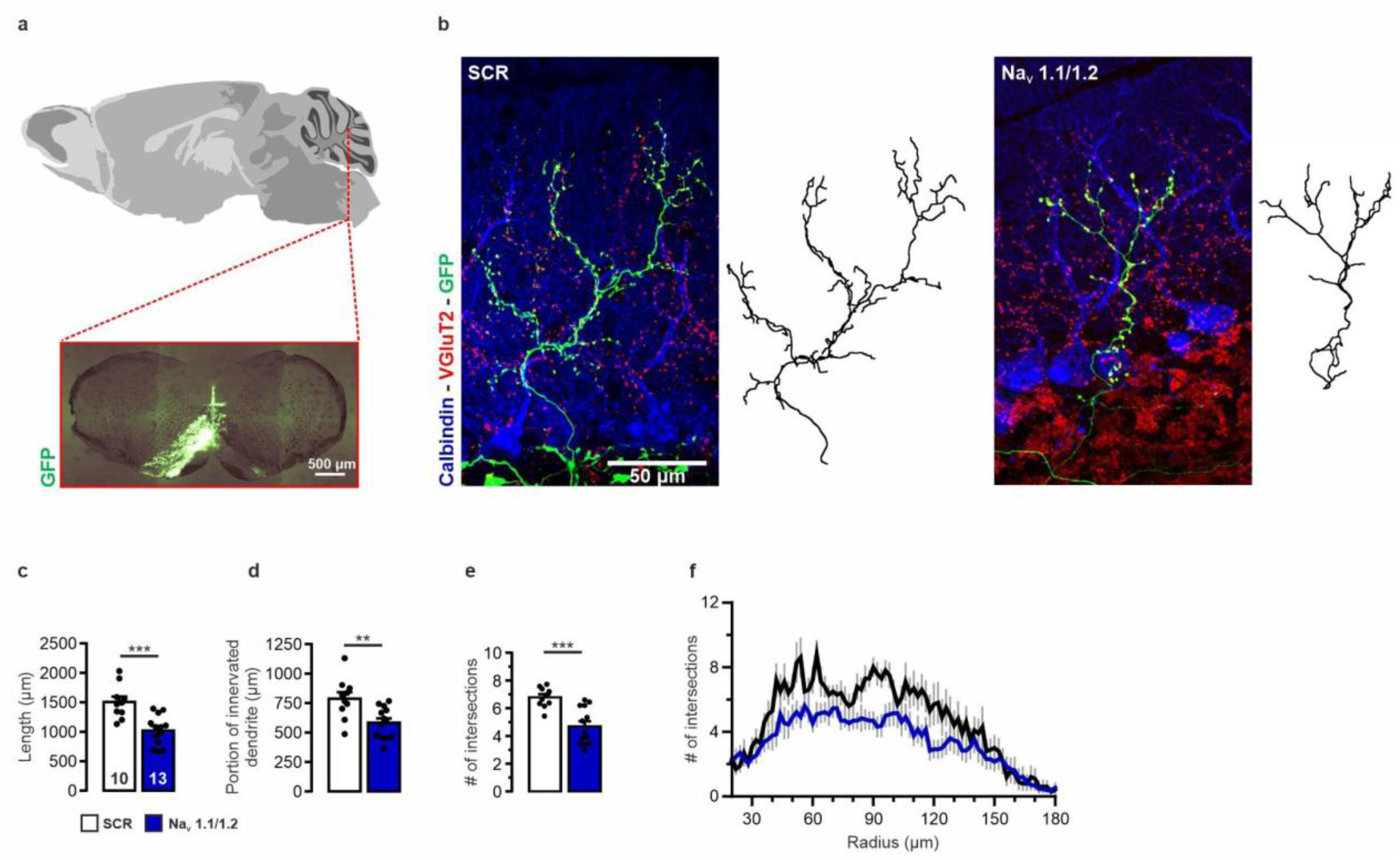
Knocking down NaV1.1/1.2 causes atrophy of CF branches. **(a)** Injection site in the IO of GFP-expressing lentiviral vectors (brainstem coronal section): GFP (green) merged with brightfield. **(b)** Representative GFP-expressing CFs (green) in sagittal cerebellar sections immunostained for VGLUT2 (varicosities, red) and calbindin (PCs, blue), and corresponding 3D reconstructed traces (black) for control (SCR, left) and NaV-KD (right) vectors. **(c-f)** NaV-KD causes a reduction of the total length of CF (**c**), of the portion of dendrite innervated by the CF (**d**), and of the number of CF branches (**e**), as assessed by Sholl analysis at 40-120 μm from the center of analysis (**f**). Scale bars: 500 µm (**a**) and 50 µm (**b**). **p<0.01; ***p<0.001.

### 4. Effect of the of NaV-KD on CF varicosities

In contrast to what observed for GAP-43-KD (see Fig. 1g-m), NaV-KD did not significantly affect the total number of varicosities per CF (NaV-KD: 169±15; control: 209±13; p=0.071; **Fig. 4a,b**) or the density of varicosities calculated over the length of innervated dendrite (NaV-KD: 2.9±0.2 varicosities/10 μm; control: 2.7±0.2 varicosities/10 μm; p=0.56; **Fig. 4d**), suggesting that the retraction of the fiber and the reduction of the number of branches could have been compensated by an increase of varicosities on the remaining branches. Indeed, NaV-KD caused an increase in the density of axonal varicosities calculated over the total length of CF branches (NaV-KD: 1.6±0.1 varicosities/10 μm; control: 1.3±0.1 varicosities/10 μm; p=0.019; **Fig. 4c**), indicating that the fewer and shorter CF branches (some of them innervating the same dendrite traits) experienced an overall increase in density of varicosities. Such an increase appears, therefore, to homeostatically compensate for the retraction caused by the NaV-KD, preserving the total number of varicosities per CF and the density of varicosities over the length of the innervated dendrite comparable to control conditions.

**Figure 4.**
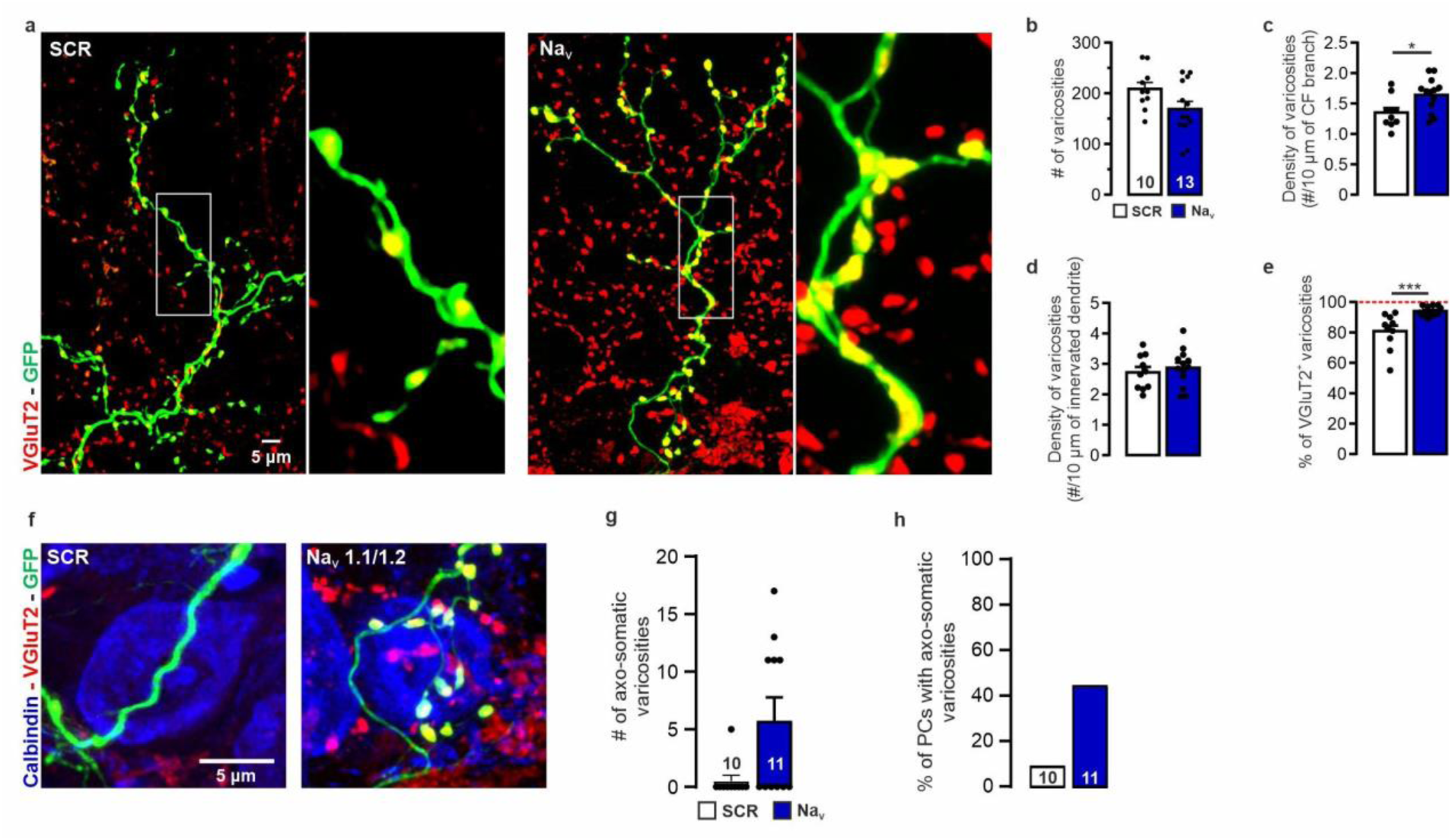
Knocking down NaV1.1/1.2 increases the density of varicosities in CFs. **(a)** Representative GFP-expressing CFs immunostained for VGLUT2 (varicosities, red; higher magnification of the images shown in Fig. 3b). **(b-e)** NaV-KD does not cause changes in the total number of varicosities (**b**), due to the increased density of varicosities on the length of the CF (**c**), resulting in unaffected density of varicosities on the length of PC innervated dendrites (**d**). **(e)** The number of VGLUT2-expressing varicosities is higher after NaV-KD. **(f)** Representative image showing PC somata (blue) innervated by GFP-positive CFs (green), and immunostained for VGLUT2 (red). **(g,h)** NaV-KD increases both the number of axo-somatic varicosities (**g**) and the frequency of PCs receiving axo-somatic synapses by CFs (**h**). Scale bars: 5 µm (**a, f**). *p<0.05; ***p<0.001.

As for GAP-43-KD, the treatment also caused an increase in the fraction of varicosities containing synaptic vesicles, identified by the presence of the vesicular glutamate transporter 2 (VGLUT2^+^ varicosities/total varicosities; NaV-KD: 94±0.8%; control: 80.8±3.71%; p=0.0009; **Fig. 4e**), suggestive of an additional compensatory mechanism in favor of functionally active presynaptic terminals.

Interestingly, NaV-KD also modified the spatial distribution of varicosities over the innervated PC: an aberrant number of somatic varicosities was present after NaV-KD (NaV-KD: 5.7±2; control: 0.50±0.50, p=0.028; **Fig. 4f,g**) with a higher occurrence of PCs bearing somatic varicosities (**Fig. 4h**). A similar trend was observed for GAP-43-KD (1.86±0.88, p=0.18; **Fig. S2**)

The transverse branches of CF, which run perpendicularly to the plane where CF and PC dendrites lie (Nishiyama et al., 2007), were previously described as the most motile branches of CF characterized by an activity-dependent control of motility, although their function is still elusive (Miyazaki et al., 2017; Nishiyama, 2014; Sugihara et al., 1999). When we analyzed the effect of NaV-KD on the CF transverse branches (**Fig. S3**), we observed a trend towards a shorter length (NaV-KD: 10.8±1.2 μm; control: 12.9±0.8 μm; number of transverse branches: 32 and 46, respectively; p=0.14), a lower number (NaV-KD: 2.5±0.7; control: 5.1±2.3; p=0.21), and a decreased density over the CF length (NaV-KD: 2.1±0.6 / mm; control: 5.1±2.3/mm; p=0.16). Although these differences were not statistically significant, they are consistent with the possibility that the reduction of CF activity may cause their destabilization and retraction. Similar trends were observed with GAP-43-KD (**Fig. S4**).

Altogether, these data show an activity-dependent control of the morphology of CF branches and density of CF varicosities. Since we observed that silencing NaV channels caused morphological modifications of CF branches overlapping with those obtained by silencing GAP-43, and since GAP-43 function can be modulated by biochemical mechanisms typically related to neuronal activity, such as binding of calmodulin and phosphorylation by protein kinase C (PKC) (Denny, 2006), it is tempting to speculate that GAP-43 mediate this form of activity-dependent control of CF morphology. To clarify whether such activity-dependent control of CF morphology was dependent on GAP-43, we analyzed GAP-43 levels after silencing of NaVs in *in vitro* primary neurons by Western blot (**Fig. S5a,b**) and *in vivo* by measuring its immunoreactivity in GFP-expressing fibers on fixed tissue (**Fig. S5c,d**). Neither method was able to detect significant differences. Although it cannot be excluded that the sensitivity of these methods is intrinsically too low, these data suggest that, GAP-43 plausible role in mediating the activity-dependent control of CF morphology occurs through different mechanisms than the modulation of GAP-43 protein level, such as phosphorylation, calmodulin binding or plasma membrane insertion (Denny, 2006; Holahan, 2017; Mosevitsky, 2005).

To rule out the possibility that the similar effects of GAP-43-KD and NaV-KD on CF branches are due to an influence of GAP-43 on neuronal excitability, we measured intrinsic excitability in GAP-43-KD primary cortical neurons. We found that GAP-43-KD, without affecting the NaV expression levels (**Fig. S6a,b**), caused a decrease in intrinsic excitability (**Fig. S6c,d**) associated with a preserved AP waveform and resting potential and a slightly decreased input resistance (**Fig. S6e-h**), consistent with GAP-43 role in supporting the formation of neuronal processes. This suggests that, while an influence on neuronal excitability by GAP-43 may not be excluded, this protein does not regulate NaV protein expression level, at least *in vitro*.

### 5. NaV-KD in CF causes an increase of dendritic spines

Previous works suggested that CF activity suppresses the formation of new spines on the dendritic territory that is innervated (Bravin et al., 1999; Cesa et al., 2007; Cesa & Strata, 2007). Therefore, we analyzed the density of spines on dendritic branches of PC innervated by NaV-KD CFs (**Fig. 5a**). We observed that spine density was increased on PC proximal dendrites of NaV-KD CFs (NaV-KD: 2.91±0.23 spines/10 μm; control: 2.02±0.015 spines/10 μm; p=0.0045; **Fig. 5b**), although no difference was observed in the fraction of spines colocalizing with varicosities or CF filaments (**Fig. 5c,d**). This effect was specific to the innervated proximal dendrite compared to distal branches, on which the spine density was not affected (Na_V_-KD: 6.43±0.78 spines/10 μm; control: 6.81±0.36 spines/10 μm; p=0.58; **Fig. 5e**).

**Figure 5.**
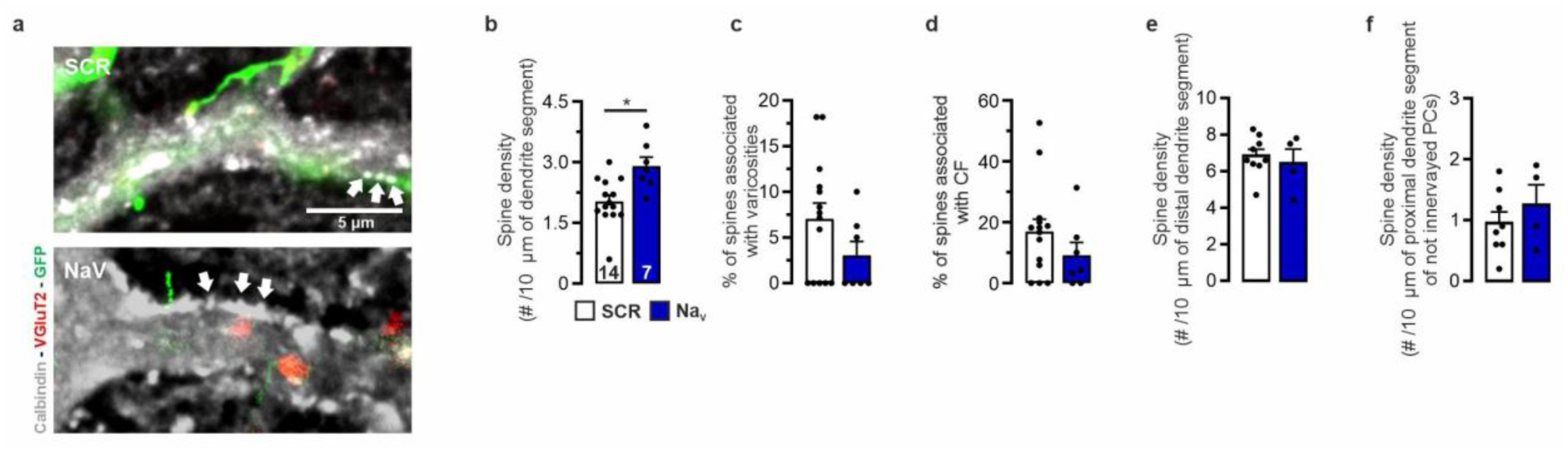
Knocking down NaV1.1/1.2 selectively increases dendritic spine density on innervated PCs. **(a)** Representative images of dendritic spines (white; arrows) on PC thick proximal branches, innervated by a GFP-positive CF (green). **(b-d)** NaV-KD causes higher dendritic spine density in the proximal segment of PCs dendrites (**b**) without affecting the percentage of spines associated with varicosities (**c**) or localized under the fiber branches (**d**). **(e,f)** The spine density assessed on distal dendritic segments of PC innervated by GFP-positive CFs (**e**) and on proximal dendritic segments of PCs not innervated by GFP-positive CFs (**f**) is not affected by NaV-KD. Scale bars: 5 µm. *p<0.05.

It is known that glutamate release from CFs activates molecular layer interneurons by spillover (Szapiro & Barbour, 2007) and that PCs innervate surrounding PCs and molecular layer interneurons by collateral axonal branches (Witter et al., 2016). Therefore, the activity of a CF and of the innervated PC may affect surrounding PCs. For this reason, we analyzed the spine density of PC neighbors of the innervated ones (within 40 μm in the same section), whose CF was GFP-negative, and observed that spine density of the proximal dendrite of these cells was unaffected (NaV-KD: 1.25±0.57 spines/10 μm; control: 0.95±0.49 spines/10 μm; p=0.41; **Fig. 5f**), demonstrating that the control of spine density exerted by single CFs only affects the innervated dendritic territory.

To assess whether the GAP-43-dependent CF remodeling is sufficient for the negative control of spine density, we analyzed the spine density of PC dendrites innervated by GAP-43 knocked-down CFs: under this condition (associated with a CF atrophy without a compensatory increase of varicosities; see Fig. 1), the resulting spine density on the innervated proximal dendrite was lower than in controls (GAP-43-KD: 1.44±0.20 spines/10 μm, p=0.027; **Fig. S7**), and not higher as in the case of NaV-KD, suggesting that CF retraction is not sufficient to induce an increase in spine density and, consequently, that the control of spine density is exerted specifically by CF activity and not by the CF trophic state.

It was previously shown that a complete blockade of electrical activity in the cerebellar cortex by intraparenchymal infusion of tetrodotoxin (TTX) caused an increase in GABAergic synapses on PC proximal dendrites (Cesa et al., 2008). However, we observed that NaV-KD in CF did not affect the density of GABAergic varicosities on PC proximal dendrites (**Fig. S8**)

Overall, these results confirm that CF electrical activity (and not just their atrophy) is able to control spinogenesis locally on the innervated dendritic territory without affecting the inhibitory connectivity.

### 6. Effect of the NaV silencing on CF-PC synaptic transmission

We finally asked whether the function of the neuronal circuit can be affected by such NaV-dependent remodeling of CF innervation (affecting its branches, varicosities, and spine density of the innervated dendritic tracts). To ascertain the functional consequences of the observed activity-dependent structural plasticity, we analyzed CF synaptic transmission by whole-cell patch-clamp electrophysiology in acute cerebellar slices. The pairing between the GFP-labelled CF and the recorded PC was individually confirmed at the end of each recording based on the paired morphology of the GFP^+^ CF and the innervated dendrite infused with a fluorescent dye in the patch pipette (Alexa Fluor 568 hydrazide; **Fig. 6a**). The amplitude of CF excitatory postsynaptic currents (EPSCs) was not significantly affected by NaV-KD (EPSC, NaV-KD: 635.1±54.0 pA, control: 538.8±54.7 pA; p=0.31; **Fig. 6b,c**). The paired-pulse ratio was slightly but significantly larger after NaV-KD (PPR at 100-ms inter-pulse interval, NaV-KD 0.88±0.01, control 0.82±0.02; p=0.048; **Fig. 6d,e**), suggesting a decrease in release probability. The EPSC had also slower kinetics, with longer onset time (NaV-KD: 3.2±0.09 ms; control: 2.03±0.29 ms; p= 0.0087) and half-width (NaV-KD: 10.2± 0.56 ms; control: 7.1± 0.7 ms; p=0.017), resulting in a higher overall synaptic charge (NaV-KD: 7325±663.2 pC; control: 4512±949.2 pC; p= 0.026; **Fig. 6f-i**). These observations therefore show that the CF structural modifications induced by NaV silencing are associated with alterations in synaptic transmission that could be relevant for the circuit function. Interestingly, similar electrophysiological results were obtained by silencing GAP-43 (EPSC, GAP-43-KD: 411±52 pA, p=0.12; PPR at 20, 50, 100, 150 and 500 ms of inter-pulse interval: treatment x interval interaction p=0.03, F (4, 56) = 2,889; onset time, GAP-43-KD: 3.3±0.4 ms, p=0.011; half-width, GAP-43-KD: 10.12±0.46 ms, p=0.0071; **Fig. S9**), suggesting the possibility that GAP-43 may mediate the observed activity-dependent plasticity of CF synaptic transmission in the NaV-KD group, consistent with its activity-dependent control and its role in synaptic transmission (Denny, 2006; Holahan, 2017; Mosevitsky, 2005).

**Figure 6.**
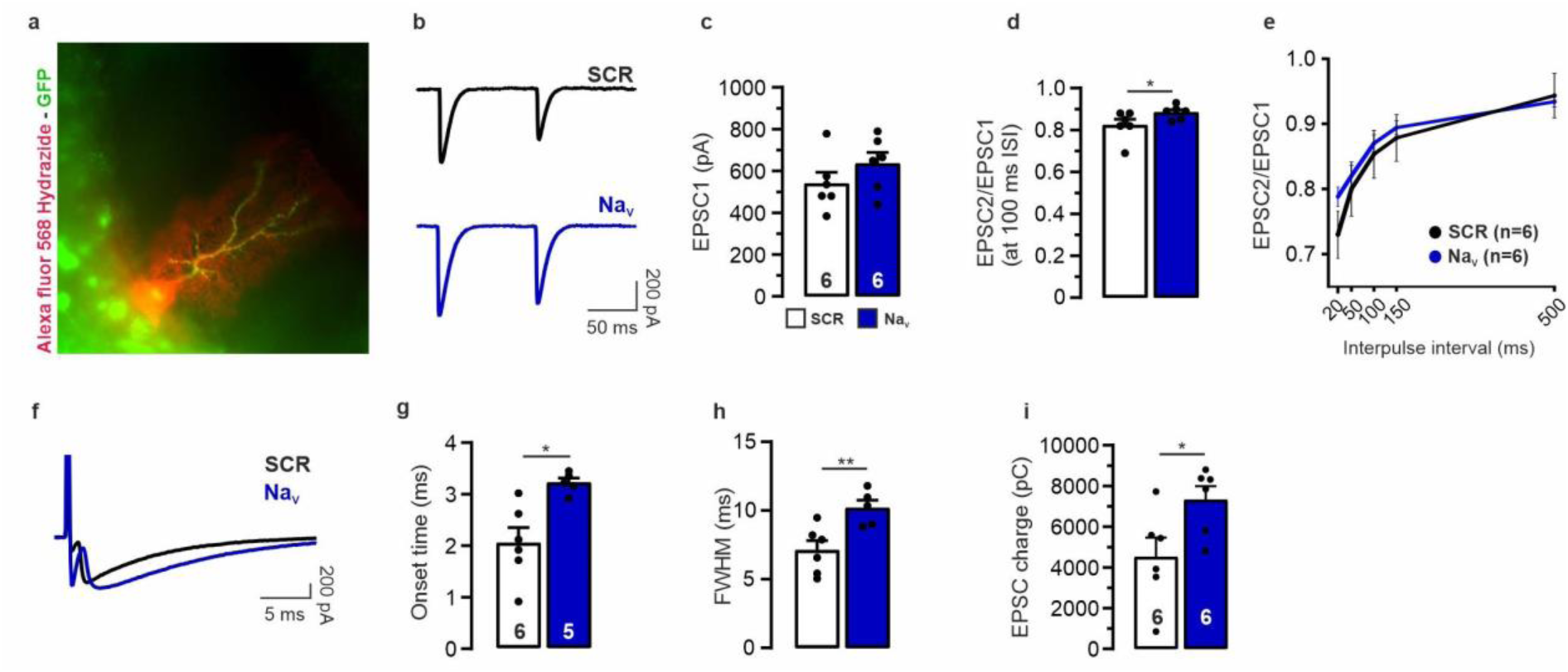
Knocking down Nav1.1/1.2 alters CF-PC synaptic transmission. **(a)** Representative GFP-positive CF paired with a patched PC, identified by pipette-infusion with red fluorescent dye. **(b)** Representative EPSC traces from NaV-KD PC innervated by CFs treated by either or control lentiviral particles. **(c)** The amplitude of CF-EPSC is not affected by NaV-KD. **(d,e)** The pair-pulse ratio, performed in the interspike interval (ISI) range 20-500 ms (**d**) is increased after NaV-KD at 100-ms ISI (**e**). **(f)** Representative CF-EPSCs recorded after either NaV-KD or SCR treatment highlight that slowdown of the EPSC kinetics after NaV-KD. **(g-i)** Analysis of the EPSC kinetics shows longer onset-to-peak time (**g**), longer half-width (**h**) and larger synaptic electric charge (**i**). Scale bars: 200 pA and 5 ms (**f**). *p<0.05; **p<0.01.

Since the integration of CF and PF synaptic inputs in PCs is crucial to determine whether long-term depression or potentiation should occur at PF synapses (Coesmans et al., 2004; Gao et al., 2012), our data show that that the activity-dependent structural plasticity of CF (possibly mediated by GAP-43) causes alterations in synaptic transmission that may affect PC function and synaptic integration.

## DISCUSSION

### Activity-dependent structural plasticity of CF main branches

Our data show that a reduction of intrinsic excitability in IO neurons, obtained by knocking-down voltage-gated sodium channels, causes a structural remodeling of cerebellar CFs, resulting in shorter and fewer branches. This effect, however, does not produce a lower number of synaptic contacts with PCs. Rather, the total number of varicosities in treated CFs is comparable to controls. This compensatory increase is also accompanied by a higher density of dendritic spines, specifically in the dendritic territory innervated by treated CFs. These data therefore demonstrate that the main branches of CFs (“ascending branches”, which innervate PCs) can undergo activity-dependent structural plasticity if the intrinsic excitability of the neurons that originate them is selectively reduced.

Previously, activity-dependent structural plasticity of CFs, triggered by harmaline-induced increase in spontaneous firing (Nishiyama et al., 2007) was observed to involve only the dynamic motility of their transverse branches, whose function is still elusive (Aimi & Yuzaki, 2023; Sugihara et al., 2000). On the other hand, remodeling of the main ascending branches was observed in reaction to lesions (Carulli et al., 2004) or after a general block of neuronal activity in the cerebellar cortex (Bravin et al., 1999). A modification of the morphology of CF varicosities was also previously observed after general block of the glutamatergic transmission (Cesa et al., 2007). These studies were based on systemic manipulations (systemic, intraparenchymal, or intraperitoneal infusion of drugs) and general blockade of neuronal/synaptic activity in the whole cerebellar area, leaving space for both homo- and heterosynaptic effects, i.e., to the possibility that complex circuit adaptations were underlying the observed effects on CF plasticity (Carulli et al., 2004). In the present study, on the contrary, we manipulated the intrinsic excitability only of a subpopulation of IO neurons to study isolated CFs with no interventions on the cerebellar cortex. Moreover, we show that the NaV-KD treatment, rather than causing a complete silencing of action potentials or NaV expression, elicits only a decrease of neuronal excitability *in vitro* and of NaV expression *in vivo*, strongly suggesting that it should be able to reduce intrinsic excitability of IO neurons without causing a complete block of CF activity, a condition which could be considered close to a physiological reduction of CF activity.

We observed that GAP-43-KD is sufficient to obtain modifications of CF branches (but not of CF varicosities) similar to those caused by NaV-KD. Since GAP-43 is a protein that stabilizes actin cytoskeleton and depends on neuronal activity and intracellular signal transduction pathways, including the interaction with calmodulin and phosphorylation by protein kinase C (Denny, 2006; Holahan, 2017; Mosevitsky, 2005), it is plausible that GAP-43 is involved in mediating the activity-dependent control of the structure of CF branches. We showed that this is unlikely to happen at the GAP-43 protein expression level, leaving the possibility that it occurs through post-translational mechanisms.

### Regulation of the density of PC spines, CF varicosities, and inhibitory inputs

Previous studies showed that PCs respond to the loss of CF input with a dramatic generation of new spines on the proximal thick dendrite, normally innervated by CF (Bravin et al., 1999; Cesa et al., 2003, 2007; Morando et al., 2001). Based on these studies, it was proposed that CF activity plays a role in locally repressing spinogenesis (Bravin et al., 1999; Cesa & Strata, 2009). Our data, by showing that CF activity negatively modulates the density of dendritic spines specifically on the innervated proximal dendrite, confirm this hypothesis. Moreover, we show that this is accompanied by an increase in the density of varicosities that fully compensates for their loss due to the shortening of the fiber branches, keeping their total number stable. Neither the increase in the density of spines nor of varicosities occurred after knocking down GAP-43. This result suggests that, while the activity-dependent structural plasticity of the brahces could be mediated by GAP-43, the regulation of the density of CF varicosities and PC spines is dependent on CF activity via different molecular mechanisms, independent of GAP-43 and the CF trophic status. We therefore speculate that changes in CF activity control on one side, CF branches (by GAP-43), and, on the other side, induce a form of homeostatic plasticity involving CF varicosities and PC spines to compensate for the decrease in CF activity.

The thick proximal PC dendritic branches are normally innervated only by CF glutamatergic inputs and GABAergic inputs originating from molecular layer interneurons (Cesa & Strata, 2009; Ichikawa et al., 2016). These inhibitory inputs can undergo activity-dependent structural and functional plasticity (Cesa et al., 2008; Scelfo et al., 2008) and are specifically reduced by the activity of PF (Ito-Ishida et al., 2014). These effects are especially relevant since structural plasticity of inhibitory inputs in the cerebellar cortex (as well as in the hippocampus) was shown to occur during learning (Ruediger et al., 2011). Moreover, it is known that the excitatory/inhibitory balance is impaired in the cerebellum under pathological conditions such as autism (Hegarty et al., 2018). Here we show that the density of inhibitory presynaptic terminals on PC proximal dendrites is, however, not directly dependent on CF activity.

### Molecular mechanisms underlying activity-dependent CF structural plasticity

As for the molecular mechanisms mediating the CF activity-dependent structural plasticity, we provide evidence supporting a possible role of GAP-43 for CF branches, consistent with our previous findings (Allegra Mascaro et al., 2013; Grasselli et al., 2011). We cannot exclude that other mediators may be involved, including soluble molecules or adhesion molecules that are released by CFs or exposed on their membrane in an activity-dependent way, such as cerebellins (CBLNs) (Yuzaki, 2011) or C1q-like family members (Kakegawa et al., 2015). In particular, C1ql1 plays a role in strengthening and preserving the dominant CF during postnatal development, by interacting with brain-specific angiogenesis inhibitor 3 (BAI3), a cell-adhesion G protein-coupled receptor that is expressed by PCs (Aimi & Yuzaki, 2023; Kakegawa et al., 2015; Moghimyfiroozabad et al., 2023; Sigoillot et al., 2015). Other soluble molecules released by CF and their receptors on PCs may also be involved in locally controlling spinogenesis, such as the corticotropin-releasing factor (CRF) (Gounko et al., 2013) and its receptors CRF-R1 (J.-B. Tian et al., 2008) and CRF-R2 (Cui et al., 2022).

### Functional implications of CF structural plasticity in cerebellar learning

We showed that the amplitude of CF EPSCs resulting from CF structural plasticity is not affected by the structural pre-/postsynaptic modifications caused by the decrease in CF excitability. We can therefore conclude that the increase in the density of spines and varicosities plays a functional compensatory role, able to keep CF synaptic transmission partially unaffected.

Although NaV-KD does not affect the CF EPSC amplitude, we observed that it causes a smaller release probability and slower current kinetics, resulting in almost double the transferred synaptic electrical charge. The observation that knocking down GAP-43 causes similar effects on current kinetics is consistent with its known role in synaptic release and vesicle mobilization, and suggests that such alterations in synaptic function are related to the activity-dependent role of GAP-43 (Denny, 2006; Holahan, 2017). The mechanisms underlying the changes in transferred synaptic charge might also be related to the homeostatic increase of CF varicosities and PC spines, involving either local accumulation of glutamate (Wadiche & Jahr, 2001) or activity-dependent modifications of Bergmann glia involved in glutamate clearance (Szapiro & Barbour, 2007). Regardless of mechanism, it is plausible that the changes in synaptic transmission caused by NaV-KD have physiologically relevant consequences: the lower release probability, longer kinetics and larger transferred charge can cause differences in PC synaptic integration, especially of trains of CF activity and with PF activity, both crucial to drive PF plasticity (Piochon et al., 2016; Titley et al., 2019). This implies, therefore, the possibility that they may ultimately affect synaptic plasticity and, consequently, cerebellar learning. Indeed, synaptic plasticity at PFs is dependent on the relative timing of CF single/multiple spikes and on the integration of intracellular calcium transients (Piochon et al., 2016; Suvrathan et al., 2016; Titley et al., 2019) and plays a crucial role in cerebellar learning (Coesmans et al., 2004; Gao et al., 2012; Ohtsuki et al., 2009). Further studies will be needed to test this hypothesis.

Previous studies have shown that, in other brain areas, including the hippocampus and the neocortex, activity-dependent changes involve the shape and number of spines and presynaptic terminals, as well as the shape of dendrites and axonal fibers, and that these changes contribute to encoding memories (Caroni et al., 2012; Holtmaat & Caroni, 2016). However, in the cerebellum, data on the contribution of structural plasticity to learning have been more limited. For instance, it was shown that learning acrobatic skills to walk on several elevated obstacles during a training lasting several weeks induces an increase in the number of PF synapses on PC in adult rats, but not of those formed by CFs or molecular layer interneurons (Black et al., 1990; Kleim et al., 1998; Stevenson et al., 2021). Based on our study, we speculate that CFs undergo experience-dependent structural plasticity involving their ascending branches, with alterations in synaptic transmission and homeostatic mechanisms that could maintain the number of their varicosities stable.

This can be related to an increase in spine density (Lee et al., 2007). This form of motor learning implies an improvement in sensory–motor integration and coordination, which occurs in the cerebellar cortex. Therefore, these observations show that learning motor skills causes an actual rewiring of the cerebellar circuit that allows a refinement of motor responses after sensory stimulations. We could reasonably expect that this includes both potentiated and depressed PF inputs (in addition to lost and newly formed synapses), and that it can result in a refined synaptic integration within single PCs, together with a redistribution of information delivered by PFs to PCs.

After eyeblink conditioning - a motor learning task used to understand the neuronal mechanisms underlying cerebellar learning - a reduction of the number of PF synapses (but not of inhibitory synapses) was found on PCs (Connor et al., 2009), resulting in disinhibition of neurons in the cerebellar nuclei. This learning task was also observed to cause outgrowth of mossy fiber collaterals in the cerebellar nuclei (Boele et al., 2013; Kleim et al., 2002; Weeks et al., 2007), which supports an enhanced response in the cerebellar nuclei to the activation of mossy fibers encoding the conditioned stimulus. Based on our data, we speculate that a detailed analysis of CF structure reveals that the structure of CF branches can be affected by this form of learning.

Other studies showed that experience-dependent structural plasticity also involves PC axon terminals (Foscarin et al., 2011) and cerebellar mossy fibers (Ruediger et al., 2011). The activity-dependent structural plasticity involving the ascending branches of CF that we have shown here represents, therefore, a potentially new piece in the puzzle of cerebellar mechanisms of plasticity contributing to encoding cerebellar memories (Gao et al., 2012). Further studies will be needed to verify this hypothesis.

### Functional implications of CF structural plasticity in disorders involving the cerebellum

Cerebellar dysfunctions contribute to the pathophysiology not only of several motor disorders, such as spinocerebellar ataxias (Robinson et al., 2020), essential tremor (Louis, 2017), multiple sclerosis (Parmar et al., 2018), and dystonia (Shakkottai et al., 2017), but also in neurodevelopmental and neuropsychiatric disorders, including autism (Wang et al., 2014) and schizophrenia (Escelsior et al., 2019). Impairments in CF connectivity to PC have been observed in several neurological and psychiatric conditions and mouse models thereof, such as in essential tremors -with a lower density of varicosities and extended innervated territory (C.-Y. Lin et al., 2014) as well as innervation of multiple PC (Kakei et al., 2021; Lang & Handforth, 2022; Pan et al., 2020; Y. C. Wu et al., 2021)- spinocerebellar ataxia type-1 (Kuo et al., 2017; Louis et al., 2023), autism (Piochon et al., 2014; Simmons et al., 2022) and schizophrenia (Veleanu et al., 2022). While it is likely that such impairments may originate, at least partially, from altered synaptogenesis or synaptic pruning during neurodevelopment, it will need to be assessed whether changes in the activity-dependent CF structural plasticity are also involved in the pathophysiology of these conditions. Interestingly, mutations in the genes SCN1A and SCN2A, encoding NaV1.1 and NaV1.2 that were knocked down in CFs in our study, are strongly associated with autism spectrum disorder and intellectual disability (Meisler et al., 2021), suggesting that the form of activity-dependent CF plasticity that we described may contribute to the pathogenesis of these diseases.

## CONCLUSIONS

Altogether, our data show that: (i) the CF electrical activity is necessary to sustain the structure of CF branches, possibly acting through GAP-43; (ii) the CF ascending branches can undergo a form of activity-dependent structural plasticity within a time window of two weeks; (iii) the density of CF varicosities and dendritic spines on target dendritic traits is affected by CF activity but through mechanisms independent of GAP-43, and they increase in response to a reduction of CF activity, as a possible homeostatic mechanism; (iv) the compensatory increase of varicosities and dendritic spines is effective in maintaining a number of varicosities on each PC and an amplitude of CF synaptic currents comparable to control; (v) the described CF structural plasticity is associated with a decreased release probability, and slower synaptic current kinetics, possibly mediated by GAP-43 and potentially affecting synaptic integration and plasticity.

In conclusion, the activity-dependent structural plasticity of CFs described here may contribute to the information encoding and implicit cerebellar learning. Further investigation will be needed to determine whether this new form of plasticity is induced by physiologically/behaviorally relevant conditions, and what role it plays in the physiology of the olivocerebellar circuit and in cerebellar learning.

## RESEARCH FUNDS

This project was supported by the European Union’s Horizon 2020 Research and Innovation Programme under the Marie Skłodowska-Curie grant agreement No 844391 (H2020 MSCA-IF-2018 “FunStructure”, GA No. 844391), by the Italian Ministry of Health (Ricerca Corrente; Ricerca Finalizzata Giovani Ricercatori GR-2019-12370176) by Ospedale Policlinico San Martino IRCCS (5x1000), and by the Italian Ministry of University and Research (EU-NextGenEU PNRR PE0000006 MNESYS)

## ACKNOWLEDGMENTS

We are grateful to Piergiorgio Strata (University of Turin, Turin, Italy), Paolo Cesare (formerly at IRCCS Santa Lucia Foundation, Rome, Italy) and Georgia Mandolesi (University of Rome San Raffaele, Rome, Italy) for scientific advice and technical assistance; Riccardo Navone (NSYN-IIT), Laura Emionite, Giuseppina Zuffanti and Michele Cilli (IRCCS San Martino) for support in animal care; Elisabetta Colombo, Diego Moruzzo, and Arta Mehilli (NSYN-IIT) for technical support; Rossana Ciancio, and Ilaria Dallorto (NSYN-IIT) for administrative support; all the other members of NSYN-IIT Center (especially Pietro Baldelli, Lorenzo Cingolani, Stefano Di Marco, Valentina Castagnola, Caterina Michetti, Letizia Zullo) for constant support and valuable scientific discussion; Christian Hansel (University of Chicago), Ilaria Carta (IIT) and Stefano Lutzu (San Martino Hospital) for valuable feedback on the manuscript.

