## Supplemental Figures for "Intrinsic excitability controls structural plasticity of cerebellar climbing fibers"

### SUPPLEMENTARY FIGURES

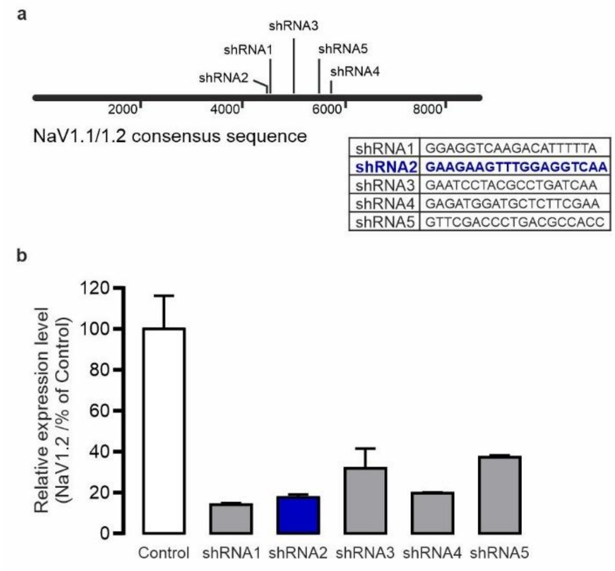


**Figure S1. Generation of lentiviral vectors**
**(a)** Five candidate shRNA sequences designed to target the identical regions of NaV1.1/NaV1.2. **(b)** Low-titer preparations tested on primary cells by real-time PCR for the expression level of NaV1.2. The shRNA2 was selected for further testing at high titer.

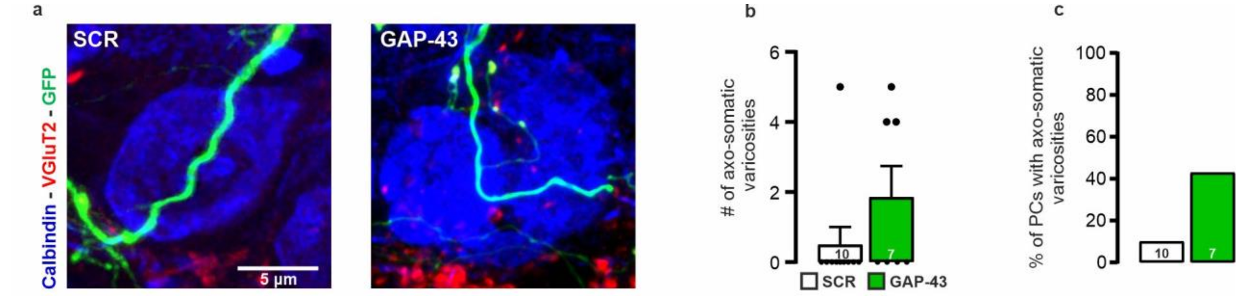


**Figure S2. The knockdown of GAP-43 in CFs does not markedly affect axosomatic varicosities**

**(a)** Somata of PCs (blue, immunostained for calbindin) innervated by GFP-positive CFs (green), immunostained for VGLUT2 (red). **(b,c)** GAP-43-KD induces a trend towards an increase in both the number of axo-somatic varicosities (**b**) and the frequency of PCs receiving axo-somatic synapses by CFs (**c**). Scale bars: 5 µm.

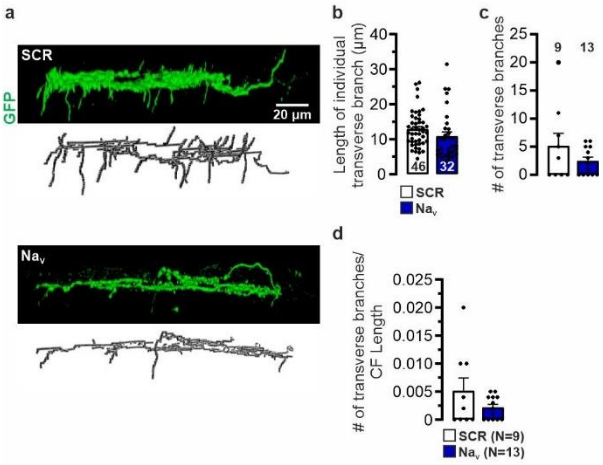


**Figure S3. The knockdown of Nav1.1/NaV1.2 does not markedly affect the transverse branches morphology of CFs**

**(a)** Lateral projection of z-stacks of GFP-positive CFs (green) acquired from sagittal cerebellar sections and corresponding 3D reconstructed traces (black) for SCR (top) and NaV-KD (bottom). Na-KD causes a trend towards a reduction of the length of individual transverse branches **(b)**, of the number of transverse branches per CF **(c),** and of their density calculated on the CF length **(d)**. Scale bars: 20 µm.

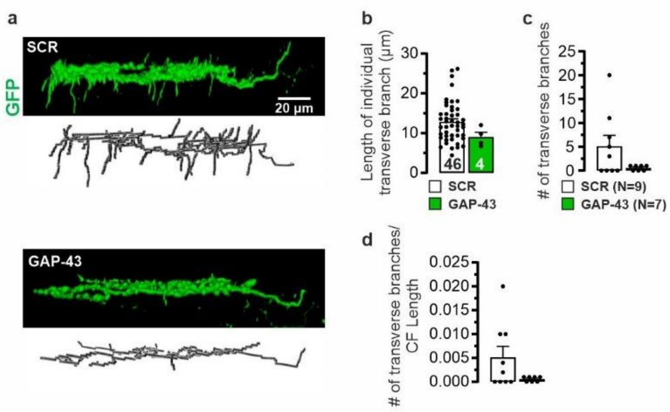


**Figure S4. The knockdown of GAP-43 does not markedly affect the transverse branches of CFs**

**(a)** Lateral projection of z-stacks of GFP-positive CFs (green) acquired from sagittal cerebellar sections and corresponding 3D reconstructed traces (in black) for SCR (*top*) and GAP-43-KD (*bottom*). GAP-43-KD does not significantly affect the average length of individual transverse branches **(b)**, their number per CF **(c),** or the number of transverse branches measured on the length of the fiber **(d)**. Scale bars: 20 µm.

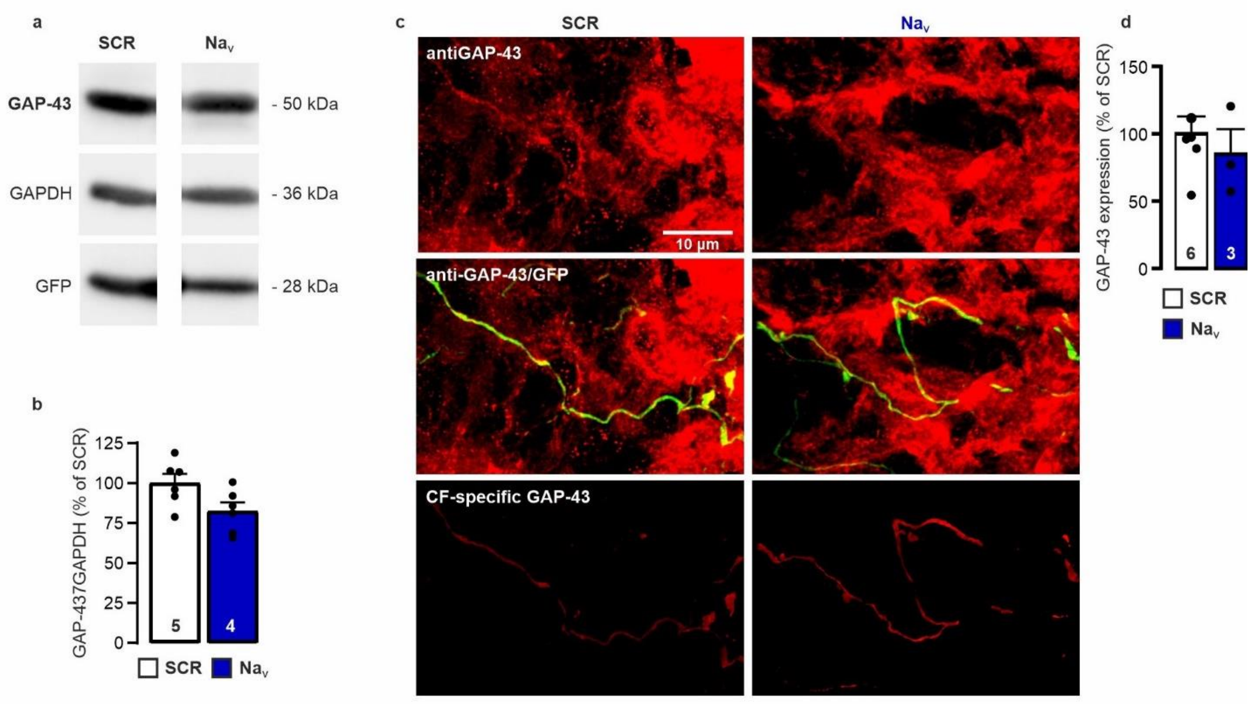


**Figure S5. The knockdown of NaV1.1/NaV1.2 does not affect the expression of GAP-43 in vitro and in vivo**

GAP-43 expression level was assessed by Western blot at days after transduction of primary cortical neurons with either SCR or NaV-KD lentiviral particles. **(a,b)** Representative immunoblot (**a**) and the densitometric quantification (**b**) shows no significant changes in GAP-43 expression.

**(c)** Z-projections of z-stacks of CFs of mice treated with either SCR (*left*) or NaV-KD (*right*) viral particles. Images corresponding to the PC layer were obtained by confocal microscopy on cerebellar slices immunostained for GAP-43 (red). Only transduced CFs are labeled by GFP (green). The fluorescent signal for GAP-43 colocalizing with GFP was isolated in each optical section and projected (lower panels). **(d)** The treatment did not significantly affect the expression of GAP-43 relative to the SCR group. Scale bars: 10 µm.

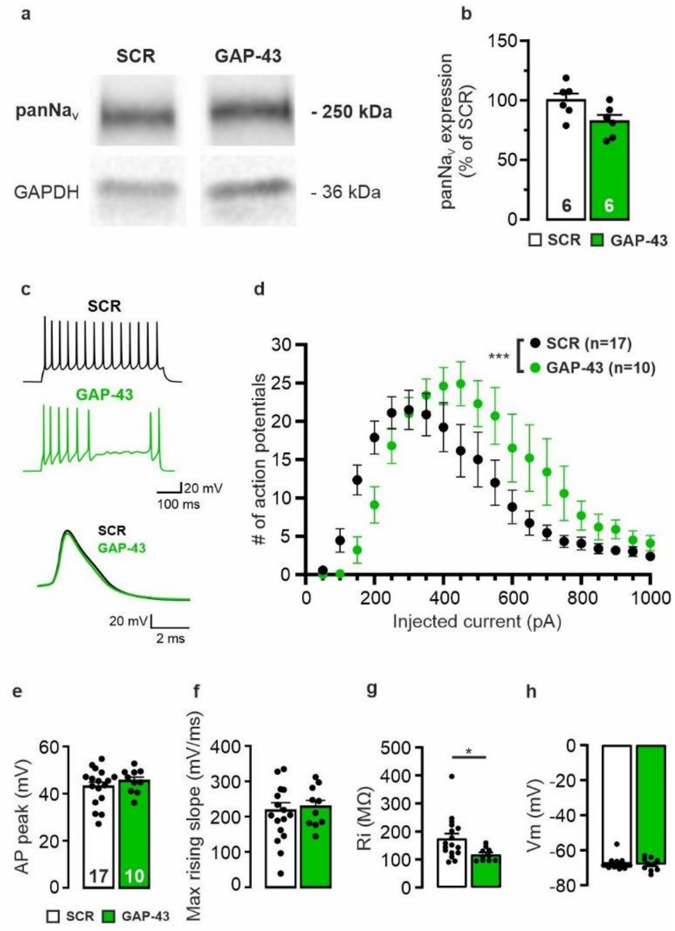


**Figure S6. The knockdown of GAP-43 decreased the intrinsic excitability in primary neurons**

**(a)** Western blotting for panNav and **(b)** quantification 7 days after transduction of primary cortical neurons (14 DIV) with either SCR or GAP-43-KD lentiviral particles (2 independent pooled experiments). **(c)** Representative traces of action potentials evoked by a depolarizing current-step (top) and corresponding first action potentials (bottom) in primary cortical neurons 7 days after lentiviral transduction (14 DIV). (d) The AP number vs injected current plot shows a decreased excitability after GAP-43-KD. **(e-h)** The analysis of the waveform of the first AP shows comparable peak potential **(e)** and maximum rising slope **(f)**. The analysis of input resistance **(g)** and membrane voltage **(h)** of recorded cells shows a significant decrease in input resistance after GAP-43-KD.

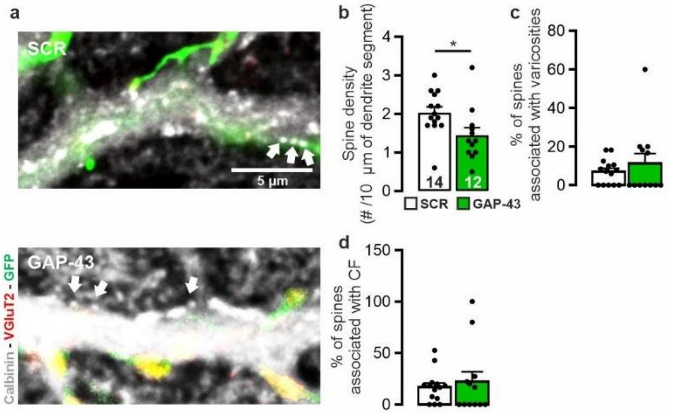


**Figure S7. The knockdown of GAP-43 decreases spine density on PC proximal dendrites**

**(a)** Dendritic spines (white arrows) on PC proximal thick branches, innervated by GFP-positive CF in SCR (*top*) and in GAP-43-KD (*bottom*). **(b)** GAP-43-KD causes a decrease in dendritic spine density in the proximal segment of PC dendrites without affecting the percentage of spines associated with varicosities **(c)** or localized under the fiber branches **(d)**. Scale bars: 5 µm. $^{*}p<0.05$.

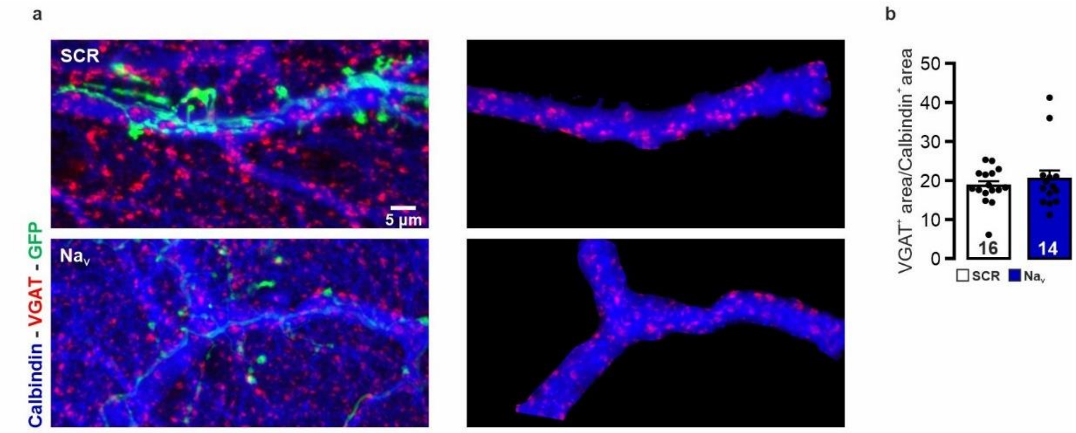


**Figure S8. The knockdown of NaV1.1/NaV1.2 does not affect the density of inhibitory inputs**

**(a)** *Left:* Inhibitory VGAT-positive varicosities (red) on PC proximal thick branches (blue, immunostained for calbindin), innervated by GFP-positive CFs (green). *Right*: inhibitory inputs impinging specifically on the trait of PC dendrite innervated by the GFP-positive CF were identified by isolating the VGAT signal colocalizing with calbindin in single optical sections, then making a z-projection of the isolated calbindin and VGAT signal. **(b)** The density of VGAT-positive varicosities specifically impinging PC branches innervated by GFP-positive CFs is not affected by NaV-KD. Scale bars: 5 µm.

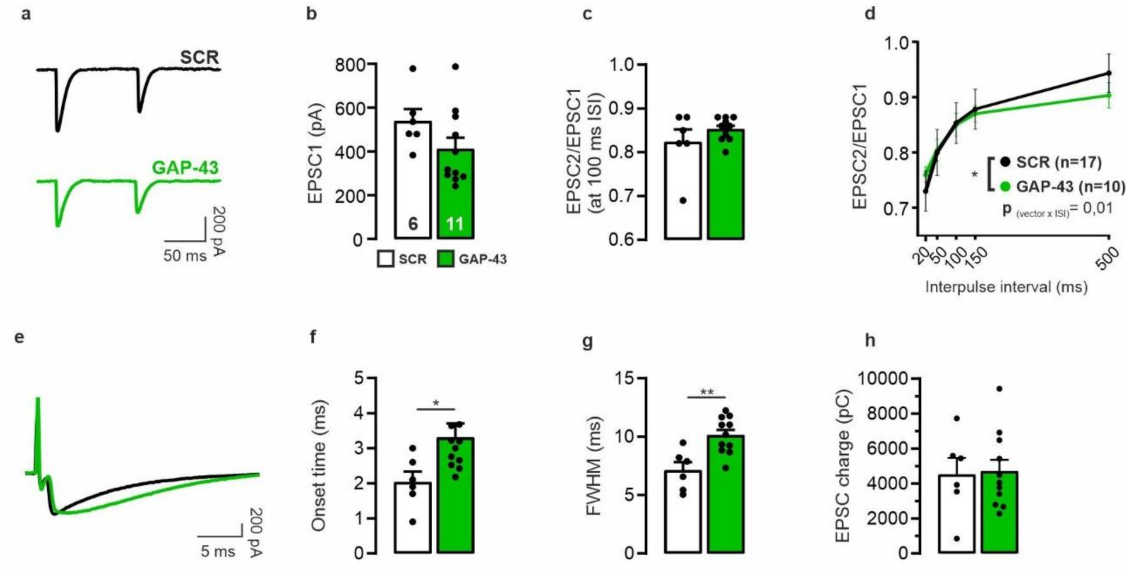


**Figure S9. Synaptic transmission after knocking down GAP-43**

**(a)** Representative EPSC traces of CFs treated with either SCR or GAP-43-KD viral vectors. **(b-d)** The CF-EPSC amplitude (**b**) and the pair-pulse ratio in the ISI range 20-500 ms (**c,d**) are not affected by GAP-43-KD. **(e-h)** The CF-EPSC kinetics are slower after GAP-43-KD (**e**), with longer onset to peak time (**f**), longer half-width (FWHM; **g**) and unchanged synaptic electric charge (**i**).
